# RNA splicing factor mutations drive myeloid neoplasm oncogenesis through protein complex poisoning

**DOI:** 10.64898/2026.09.29.754980

**Authors:** Pedro L Moura, Rui M Branca, Yaroslav Kaminskiy, Ioannis Siavelis, Kovi R. Shrung, Sophia Hofmann, Christina M Perraki, Alexandra Argyriou, Sadaf Fazeli, Masahiro M Nakagawa, Teresa Mortera-Blanco, Steffen Boettcher, Caroline Kubaczka, Thorsten M. Schlaeger, Maria Creignou, Indira Barbosa, Ann-Charlotte Björklund, Mikaela Hillberg Widfeldt, Dennis Bosch, Sanne Massaar, Mathijs Sanders, Fredrik Fagerström-Billai, R. Grant Rowe, Petter S Woll, Sten Eirik W Jacobsen, Johanna Ungerstedt, Yasuhito Nannya, Vanessa Lundin, Seishi Ogawa, Janne Lehtiö, Eva Hellström-Lindberg

## Abstract

RNA splicing factor mutations (SF^mut^) are founding oncogenic events which cause RNA splicing errors with unpredictable gene expression dynamics. Despite extensive transcriptomic studies, SF^mut^ cancer-initiating mechanisms remain elusive. Among SF^mut^ cancers, myeloid neoplasms (MN) alone offer a setting where true cancer-driving SF^mut^ stem cells can be identified, namely the hematopoietic stem cell (HSC). Within a cohort of 62 MN patients and 20 healthy donors, we conducted long/short-read single-cell transcriptomics (10X-ONT, *n* = 21) and immunophenotype-resolved low-cell proteomics (*pauciproteomics, n* = 78) to resolve gene expression, RNA splicing and protein expression dynamics across healthy and SF^mut^ MN hematopoiesis. During hematopoietic differentiation, SF^mut^ RNA mis-splicing decouples proteotranscriptomic dynamics in a mutation-specific manner. Pauciproteomics defines the functional effects of SF^mut^ RNA mis-splicing on the protein layer, identifying early-stage protein effects which poison entire functional protein complexes and progressively disrupt the proteome-wide network during cellular maturation. Importantly, this information could neither be resolved nor predicted by transcriptomic analyses. Finally, we functionally validate the proteome dynamics of HSC through induced pluripotent stem cell culture models of early hematopoietic differentiation, identifying candidate mechanisms for SF^mut^ oncogenesis. Overall, these data elucidate SF^mut^ MN disease biology with unprecedented granularity, defining the molecular consequences of SF^mut^ RNA mis-splicing and offering a generalizable framework for cancer stem cell studies.

## Introduction

Genetic aberrations in oncogenic driver genes are associated with cancer establishment and progression.^1^ Many cancer driver mutations affecting key processes are well-explored in terms of their effects on RNA and protein expression, as well as their functional consequences for cell behavior. Identifying such mechanisms has served as a basis for breakthroughs in therapeutic precision, vastly improving outcomes for molecularly defined patient groups.^2–5^

The spliceosome and its interactors are central for RNA splicing, mature mRNA formation, and correct protein synthesis. Two pivotal publications in 2011 unexpectedly revealed driving and co-driving RNA splicing factor mutations (SF^mut^) in ∼50% of myeloid neoplasms (MN), including myelodysplastic syndromes (MDS), myeloproliferative neoplasms (MPN) and mixed MDS-MPN.^6,7^ These mutations lead to RNA splicing errors (mis-splicing), which frequently cause nonsense-mediated RNA decay (NMD), and have been linked to aberrant cellular growth and differentiation, as well as a remarkable clonal advantage in malignancy. Hematopoietic stem cell (HSC)-borne mutations in the *SF3B1* gene can drive MN on their own, enabling a decades-long process of clonal expansion with infrequent co-mutations.^8–10^ Mutations affecting the *SRSF2*, *U2AF1* and *ZRSR2* genes have also been identified as key risk factors for MN development in clonal hematopoiesis (CH), again justifying an HSC origin and long-term clonal expansion process for these mutations^11,12^. This differs from other cancers, wherein SF^mut^ occur as a subclone which gives rise to a rapidly progressive disease (*e*.*g*. breast cancer, lung adenocarcinoma, uveal melanoma). Importantly, irrespective of clonal or subclonal establishment, SF^mut^ are enigmatic cancer drivers with unexplained cancer-driving mechanisms and lacking therapeutic avenues. Considering that SF^mut^ drive ∼350,000 new cases per year worldwide in MN alone, this is an unmet medical need of large proportions.^13^

Contrasting with the clear expansion ability of SF^mut^ cancer cells in the human body, genetic and cellular models of SF^mut^ display significant growth disadvantages, for reasons yet to be understood.^14–17^ As a starting point for clonal expansion in CH and MN, the HSC provides the focal point to resolve the role of these mutations in cancer. The obvious limitations to such studies are that 1) HSCs are sparse within the bone marrow (<0.3% of cells), creating a technical challenge for their study; and 2) despite advances in biological models, no *in vitro* method can preserve HSC stemness alongside cell expansion. Thus, while we know the SF^mut^ HSC is a cancer stem cell in MN, their cancer-promoting biology remains poorly defined.

In this study, we meet this need by developing a highly sensitive workflow to resolve RNA splicing, RNA expression and protein level dynamics in rare cell compartments. Specifically, we couple the 10X Genomics platform with hybrid Illumina-Nanopore sequencing to acquire dual short/long-read single-cell transcriptomes (*10X-ONT*), alongside a sensitive mass spectrometry-based workflow for immunophenotype-resolved proteomics, generalizable to the analysis of any low-cell sample (termed *pauciproteomics*). Using these technologies, we define the RNA and protein dynamics of SF^mut^ early hematopoiesis, identifying RNA mis-splicing as the seed for gradual and catastrophic dysregulation of the overall protein network. Analysis of the HSC compartment defines candidate oncogenic mechanisms, which we verified recur in induced pluripotent stem cell models. Overall, these data elucidate SF^mut^ MN disease biology with unprecedented granularity, defining the molecular consequences of SF^mut^ RNA mis-splicing and offering a generalizable framework for cancer stem cell studies.

## Results

### RNA mis-splicing decouples transcriptomic and proteomic dynamics in SF^mut^ MN

The proteotranscriptomic analysis of adult human bone marrow HSCs precludes the use of conventional methods due to their scarcity (< 0.3% of all bone marrow cells^18^). In a previous study, we pursued plate-based genotyping and single-cell RNA sequencing (scRNAseq) of HSCs and showed that HSCs are significantly affected by mutation-induced RNA mis-splicing.^9^ However, this plate-based approach is difficult to scale up for a broader unbiased analysis of differentiation dynamics within a larger patient population. Conversely, from a proteomics perspective, single-cell proteomics has been successful to define highly expressed proteins across HSCs and other cell compartments,^19^ but its limited sensitivity and scalability constrain its use for larger cohorts. Notably, the cited pivotal study investigating single-cell proteomic hematopoiesis included only two individuals. Therefore, the key open challenges yet unmet in the literature are to 1) precisely define RNA expression and splicing dynamics in HSC and during downstream hematopoietic differentiation; 2) globally profile the HSC proteome with maximal resolution; and 3) assess a large clinical cohort to derive definitive, disease-representative information.

To meet these objectives, we pursued an in-depth single-cell transcriptomic analysis (scRNAseq) of gene expression and RNA isoform dynamics by sequencing 10X Chromium-derived cDNA with both Illumina short-read and Oxford Nanopore Technologies (ONT) PromethION long-read RNA sequencing and 10X Chromium (10X-ONT). Concurrently, to combine the high sensitivity of conventional proteomics with cell compartment specification, we devised pauciproteomics, a mass spectrometric approach based on the purification of immunophenotypically-defined cell compartments followed by automated sample handling and analysis in a mass spectrometer with single-cell analysis capacity. Importantly, these two technologies make analysis of reasonably large cohorts feasible, enabling us to confidently profile RNA and protein dynamics in SF^mut^ MN at the stem cell level (**Fig. 1A, Data S1**).

**Figure 1:**
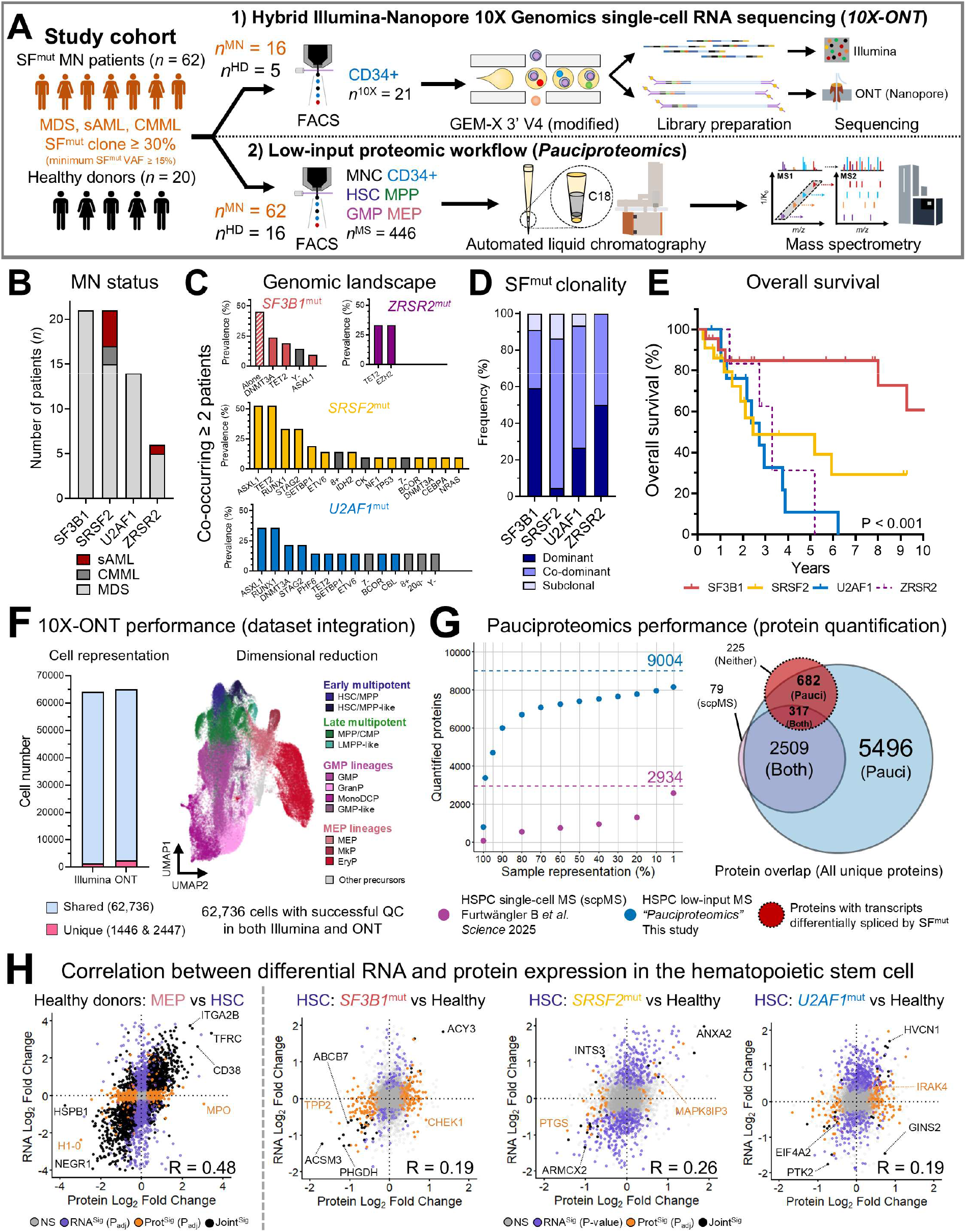
RNA splicing factor mutations decouple proteotranscriptomic dynamics in hematopoietic stem cells. **A)** Summary workflow for integrated analysis of RNA expression and splicing dynamics at single-cell resolution (*10X-ONT*, single-cell compartmentalization with the 10X Genomics platform followed by parallel Illumina short-read RNA sequencing and Oxford Nanopore Technologies [ONT] long-read RNA sequencing), coupled with an optimized workflow for high-throughput, low-cell mass spectrometry analysis (*Pauciproteomics*). **B)** Distribution of SF^mut^ myeloid neoplasm (MN) diagnoses in the study cohort (myelodysplastic syndrome [MDS], chronic myelomonocytic leukemia [CMML], secondary acute myeloid leukemia [sAML]). **C)** Mutational and cytogenetic abnormality frequencies in each major SF^mut^ category (only shown if present in two or more patients), “Alone” indicates cases with isolated SF^mut^. A complete oncoplot is provided in **Figure S1A**. **D)** SF^mut^ clonality state per SF^mut^ category, as determined by variant allele frequencies of all identified driver mutations. **E)** Kaplan-Meier plot of overall survival per SF^mut^ category. P-value corresponds to an analysis of variance (ANOVA) across all SFmut categories except for *ZRSR2*^mut^ (excluded due to low numbers). **F)** Cell numbers obtained per each sequencing technology after successful quality control steps (left; light-blue corresponds to cells obtained in both technologies, pink corresponds to cells obtained only in one technology) and UMAP-based bidimensional projection of cells captured by both technologies, labelled and colored accordingly to the major myeloid progenitor lineages (right). Sample quality control details in **Figure S2**. **G)** Quantitative performance of this study’s pauciproteomics dataset (blue) as compared to state-of-the-art single-cell proteomics of the HSPC compartment (Furtwängler B *et al*., *Science* 2025, magenta) (left). The Venn diagram depicts the overlap of protein detection between both studies, and the group of proteins with transcripts characterized to be mis-spliced (red). Experiment quality control details in **Figure S3**. **H)** Proteotranscriptomic comparison of relative protein (X-axis) and RNA (Y-axis) abundance within cell lineages in a normal background (left-most panel, MEP vs HSC), in contrast to HSC proteotranscriptomes in each SF^mut^ category *vs*. healthy donor HSCs (from left to right, *SF3B1*^mut^, *SRSF2*^mut^, *U2AF1*^mut^). Correlation coefficients (R) are indicated in the bottom-right corner of each graph. Differential expression test significance is annotated by color (black = significant in both RNA and protein, Joint^Sig^; orange = significant in protein but not in RNA, Prot^Sig^; purple = significant in RNA but not in protein, RNA^Sig^; grey = non-significant in either dataset, NS). Significance cut-offs RNA: *p* < 0.10, |Log2FC| > 0.25; Pauciproteomics: FDR < 0.10, |Log2FC| > 0.25. **Cell compartment legend: HSPC:** Hematopoietic stem and progenitor cell. **HSC**: Hematopoietic stem cell. **MPP**: Multipotent progenitor. **CMP**: Common Myeloid Progenitor. **GMP**: Granulocyte-Monocyte Progenitor. **MEP**: Megakaryocyte-Erythroid Progenitor. **GranP**: Granulocyte-committed progenitor. **MonoDCP**: Monocyte/Dendritic cell-committed progenitor. **MkP**: Megakaryocyte-committed progenitor. **EryP**: Erythroid-committed progenitor.

Our MN study cohort was based on defined inclusion criteria and sample availability (**Methods**). The diagnoses considered were SF^mut^-driven MDS, chronic myelomonocytic leukemia (CMML) and secondary acute myeloid leukemia (sAML), totaling 62 MN cases (SF^mut^ variant allele frequency ≥ 15% as determined by NGS panels), together with a comparative cohort of healthy SF^wt^ BM donors (*n* = 20). Overall, the distribution of SF^mut^ patients was well-balanced between *SF3B1*^mut^, *SRSF2*^mut^ and *U2AF1*^mut^ cases with, as expected, fewer *ZRSR2*^mut^ cases (**Fig. 1B**). This cohort was typical in terms of genetic/cytogenetic profiles (**Fig. 1C**, **Fig. S1A**) and clinical manifestations (**Fig. S1B**).^7,20,21^ SF^mut^ were the dominant or co-dominant genetic aberration in most cases, supporting the HSC origin and long-term expansion of SF^mut^ clones in these patients (**Fig. 1D**). The risk profile within this cohort was similarly typical for each SF^mut^ subgroup (**Fig. 1E**).

10X-ONT was utilized for analysis of total CD34^+^ hematopoietic stem and progenitor cells (HSPC) in 21 of the 82 individuals (**Fig. S2A, Data S2**). After successful quality control and sample demultiplexing (**Fig. S2B-S2F**), 62,726 cells were obtained with matching data in both short- and long-read RNA sequencing and recapitulating the major myeloid lineages (**Fig. 1F, Data S3**). A few MN cases displayed malignant gene expression of specific cell compartments; in all instances, these cells retained markers of the original compartment and were labelled accordingly (“HSC/MPP-like”, “LMPP-like”, “GMP-like”, Fig. **S2E/F**).

Pauciproteomics was pursued for 78 of the 82 individuals, focusing on immunophenotypically defined HSCs, multipotent progenitors (MPP), granulocyte-monocyte progenitors (GMP) and megakaryocyte-erythroid progenitors (MEP), together with the clinically relevant but heterogeneous compartments of CD34^+^ HSPC and BM MNC, totaling 446 samples (**Fig. S3A-C, Data S2**). The capacity of pauciproteomics for protein detection and quantification was robust, tripling the maximum depth of single-cell proteomics (**Fig. 1G**, **Fig. S3D**, median of 7400 proteins quantified; and in total, 9004 proteins were quantified from 220,713 precursors and 159,328 peptides), and enabling in-depth proteomic studies, including digital sample sorting (**Fig. S3E**), quantification of low-abundant proteins (**Fig. S3F**) and proteome-based mapping of cytogenetic aberrations (**Fig. S3G**).

Next, we integrated the resulting data from both 10X-ONT and pauciproteomics to study proteotranscriptomic dynamics within the HSC compartment (**Fig. 1H, Data S4**). Comparing healthy HSC to other hematopoietic lineages displayed the typical pattern of partial correlation between RNA and protein moieties (*e.g*. HSC vs MEP *R*=0.48). In stark contrast, the differential expression patterns imparted by SF^mut^ in HSC were associated with a striking lack of RNA/protein correlation (*SF3B1*^mut^ *R*=0.19, *SRSF2*^mut^ *R*=0.26, *U2AF1*^mut^ *R*=0.19).

### Proteotranscriptomics defines the global consequences of SF^mut^ RNA mis-splicing

We hypothesized that this lack of RNA/protein correlation would be primarily caused by RNA mis-splicing. Mapping alternative RNA splicing events (ASEs) in 10X-ONT data displayed ASE clustering as a clear characteristic of the original SF^mut^, including separation of *U2AF1*^Q157^ and *U2AF1*^S^^34^ hotspots (**Fig. 2A**), due to which the following analyses were pursued in a mutation-specific manner. For analysis of splicing dynamics, ASEs were mapped across cell differentiation pseudotime, as calculated by the Monocle3 algorithm (**Fig. 2B**). Within the *SF3B1*^mut^ setting, mis-splicing was largely stable from the HSC point forward (**Fig. 2C**). Thus, coupling these ASE data with a comprehensive literature database (**Data S5**) and further validation in previous SF^mut^ MNC/CD34^+^ RNA sequencing data from our constellation,^22^ we assessed whether ASE effects on the proteome were similarly stable over differentiation.

**Figure 2:**
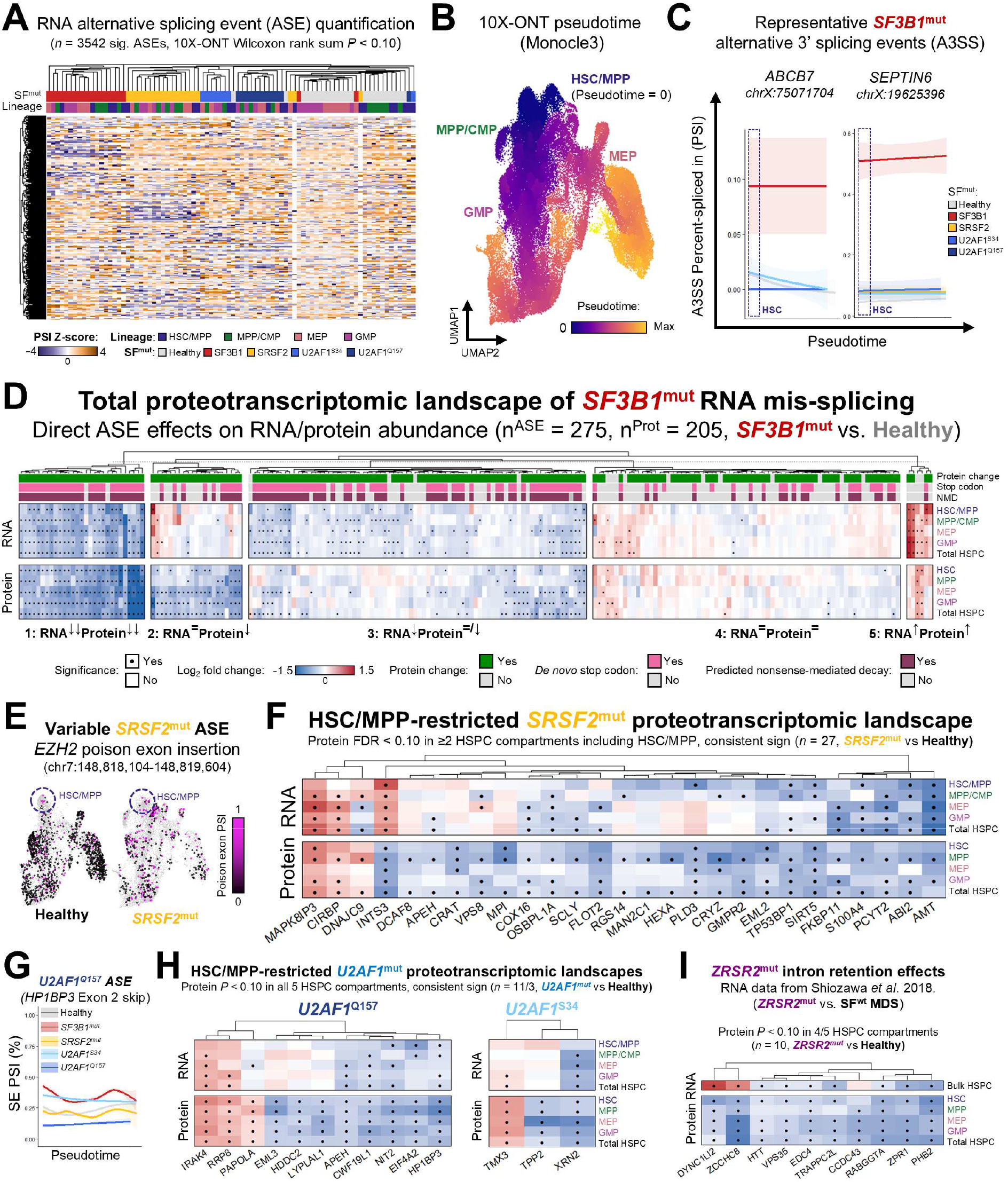
Disturbed RNA splicing remodels the protein landscape throughout early SF^mut^ hematopoiesis. **A)** Heatmap of percent spliced-in [PSI] Z-score values from significantly alternatively spliced junctions (alternative splicing events [ASEs]) of hematopoietic stem and progenitor cells quantified in 10X-ONT data. To minimize ASE dropout, single-cell data were pseudobulked into major lineage cell compartments for each sample (HSC/MPP, MPP/CMP, MEP, GMP). Mutation metadata is labelled above the heatmap. Unsupervised hierarchical clustering of sample/lineage pseudobulks was performed using the complete linkage method based on Euclidean distances. PSI Z-scores are labelled from purple (cut-off -4) to orange (cut-off +4), where Z-score indicates predominance of alternative junction usage. Columns correspond to sample/lineage pseudobulks, rows correspond to individual ASEs. **B)** UMAP visualization of cell distance-based pseudotime values, obtained through Monocle3. The HSC/MPP and HSC/MPP-like compartments were defined as starting points for pseudotime calculation, and cell distances were otherwise calculated in an unbiased manner. **C)** Smoothed mean PSI values of significantly alternatively spliced *SF3B1*^mut^-specific junctions mapped against pseudotime, starting at pseudotime = 0 and progressing to maximum pseudotime values in the data. PSI curves were modelled using a Generalized Additive Model (GAM) with regularized smoothing splines and separated by mutation. **D)** Relative RNA/protein heatmap (*SF3B1*^mut^ vs. Healthy) of genes with significant *SF3B1*^mut^ ASEs. Predicted ASE effects on the protein sequence, stop codon induction and nonsense-mediated decay (NMD) dynamics are marked above the heatmap. RNA/protein clusters were obtained by unsupervised K-means clustering (K = 5) and labelled with the predominant direction of RNA/protein expression within the cluster. Columns correspond to individual genes, rows correspond to cell compartments analyzed in pseudobulk 10X-ONT (RNA) or pauciproteomics (Protein) compared between mutant and healthy cells. Log_2_ fold change (Log_2_FC) values are colored from blue (negative Log_2_FC, lower expression in mutant cells) to red (positive Log_2_FC, higher expression in mutant cells). Dots indicate differential expression significance in a given cell compartment (RNA: *p* < 0.10, |Log2FC| > 0.25; Protein: FDR < 0.10, |Log2FC| > 0.25). **E)** UMAP visualization of *EZH2* alternative transcript splicing in healthy (left) and *SRSF2*_mut_ (right) conditions. **F)** Relative RNA/protein heatmap (*SRSF2*^mut^ vs. Healthy) of genes whose encoded proteins are significantly differentially expressed within the HSC and/or MPP compartments and are associated with significant *SRSF2*^mut^ ASEs. **G)** Smoothed mean PSI of an exon skipping (SE) ASE affecting exon 2 of the *HP1BP3* gene. **H/I)** Relative RNA/protein heatmaps of proteins differentially expressed in all five assessed HSPC compartments and associated with **H)** significant ASEs in *U2AF1*^Q157^ and *U2AF1*^S^^34^ conditions and **I)** significant intron retention events in *ZRSR2*^mut^ conditions.

Overall, transcriptomic information alone failed to predict downstream ASE consequences, despite clear discrimination of cryptic splicing site locations and prediction of NMD events. Indeed, most *SF3B1*^mut^ ASEs induced *de novo* stop codons at locations expected to induce NMD, and therefore NMD-associated transcript degradation. Yet, only a small subset of these ASEs significantly reduced both RNA and protein levels (cluster 1), while many ASEs induced only protein deficiency (cluster 2), only RNA deficiency (cluster 3) or no change whatsoever (cluster 4) despite possessing the same features (**Fig. 2D**, **Fig. S4A**). Interestingly, most protein alterations were introduced already at the HSC stage and continued with the same magnitude and direction during differentiation. ASEs associated with highly significant downregulation in both RNA and protein included known *SF3B1*^mut^ RNA mis-splicing targets (*e.g. ABCB7*, *RDX*, *UBA7*, *MAP3K7*), but most other ASE-associated protein effects were hitherto uncharacterized, as they could not be predicted by transcriptomics alone.

Further, 10X-ONT long-read data showed to be essential in order to dissect intricate transcript dynamics; for example, mis-splicing of *DLST* transcripts produced not only a cryptic isoform with one alternative 3’ splicing event linked to NMD, but also a “doubly mis-spliced” transcript where an exon skipping event upstream of the alternative 3’ splicing event caused the novel transcript to retain a functional open reading frame through its entirety (**Fig. S4B**).

Pursuing the same analysis in the *SRSF2*^mut^ setting defined a much larger number of ASEs and yet significantly lower proportions of RNA or protein alterations as compared to the *SF3B1*^mut^ setting. We hypothesized this may be associated with *SRSF2*^mut^ alternative exon skipping being less likely to introduce major transcript disruption versus *SF3B1*^mut^ cryptic 3’ splicing^23–26^ (**Fig. S5A**). Therefore, we restricted the detailed analysis of *SRSF2*^mut^ ASEs to only include proteins with concerted protein expression changes across distinct HSPC compartments (**Fig. S5B**).

Unlike *SF3B1*^mut^, and except for few proteins already disturbed in HSC and/or MPP, most *SRSF2*^mut^ ASE-associated proteomic effects were restricted to the GMP compartment as well as the broader CD34 compartment, which in *SRSF2*^mut^ is skewed for GMP predominance (**Fig. S3B**). This was also recapitulated within the RNA splicing layer; for example, relatively normal *EZH2* RNA splicing, RNA expression and protein levels in HSC/MPP only give rise to *EZH2* mis-splicing and deficiency upon GMP/MEP differentiation (**Fig. 2E, Fig. S5C-E**). In fact, only 25% of all proteomic effect-associated *SRSF2*^mut^ ASEs displayed any significant protein shift in HSC or MPP, of which nearly half did not induce RNA expression alterations (**Fig. 2F**). However, proteins altered within these restricted sets of ASEs correlated very well with the same proteins in external *SRSF2*^mut^ AML proteomic cohorts, despite low correlation both between the two external cohorts and between pauciproteomics and each cohort (**Fig. S5F**). Importantly, since *SRSF2*^mut^ ASEs do modulate the HSC proteome, these data support a role for *SRSF2*^mut^ in long-term clonal establishment and/or evolution processes within HSC.

Next, we assessed *U2AF1*^mut^ cases. These were grouped by hotspot for proteomic analysis, since 10X-ONT data displayed significant hotspot specificity of *U2AF1*^mut^ ASEs, (**Fig. 2A**, *e.g. HP1BP3* ASE in **Fig. 2G**). Due to the hotspot split, and coupled with relatively lower sample numbers, this analysis was statistically less powered than either the *SF3B1*^mut^ or *SRSF2*^mut^ analyses. Nonetheless, *U2AF1*^mut^ hotspot-specific ASEs were again specifically associated with significant proteome alterations consistent from HSC and onward, including known protein events IRAK4 and EIF4A2 (**Fig. 2H**).

Finally, although lacking 10X-ONT data, we investigated the *ZRSR2*^mut^ proteomic phenotype and identified a clear reduction in protein level for genes affected by *ZRSR2*^mut^ intron retention events (**Fig. 2I**, **Fig. S6A/B**), as well as for specific proteins that interact with these same gene products, yet not themselves affected by RNA mis-splicing. This is exemplified by the prohibitin (PHB) complex, formed by PHB1 and PHB2, where *PHB2* alone displays significant RNA mis-splicing in bulk transcriptomics data of magnetically purified CD34^+^ HSPCs^22^ (**Fig. S6C**), and yet both proteins were decreased in *ZRSR2*^mut^ samples (**Fig. S6D**).

In summary, SF^mut^ ASEs directly disrupt protein abundance during hematopoietic cell differentiation of all four SF^mut^ groups. This direct influence of SF^mut^ ASE on the proteome extends to significant and widespread proteomic disruption of the HSC compartment. Thus, pauciproteomics defines the functional protein layer that ultimately links SF^mut^ RNA mis-splicing to clonal HSC expansion and, in turn, to MN development.

### Aberrant SF^mut^ RNA splicing gradually poisons the proteome-wide network during hematopoietic differentiation

SF^mut^ RNA mis-splicing has been mechanistically linked to protein complex disruption with clinical relevance. For example, *SRSF2*^mut^-induced alternative RNA splicing was shown to drive *INTS3*-mediated loss of the integrator complex^23^ and *EZH2*-mediated loss of the EZH2/EED complex.^27^ Given the high resolution achieved by 10X-ONT and pauciproteomics on dissecting the direct impact of individual RNA mis-splicing events, we next extended our analysis to a global overview of disturbed RNA and protein dynamics within the *SF3B1*^mut^ and *SRSF2*^mut^ settings. To do so, we utilized the STRING protein-protein interaction database^28^ as a scaffold protein network and identified clusters seeded by all differentially expressed genes at RNA and protein level within any of the hematopoietic lineages (FDR < 0.10 and |Log_2_FC| ≥ 0.25 within each compartment, **Data S6**), since protein-protein interactions may vary with cell identity and yet retain similar overarching functional roles.

Strikingly, the functional patterns identified within RNA and protein data were completely distinct in the *SF3B1*^mut^ setting. RNA data were nearly devoid of functional meaning despite a substantial number of differentially expressed genes (*n*^DEG^ = 91), many of which ASE-driven. In contrast, protein data displayed significant enrichment of numerous protein clusters with functional roles (*n*^DEP^ = 717) and largely associated with *SF3B1*^mut^ ASE (**Fig. 3A, Fig. S7**). This is further highlighted by the much higher connectivity of differentially expressed genes in protein data as compared to RNA data (**Fig. 3B**).

**Figure 3:**
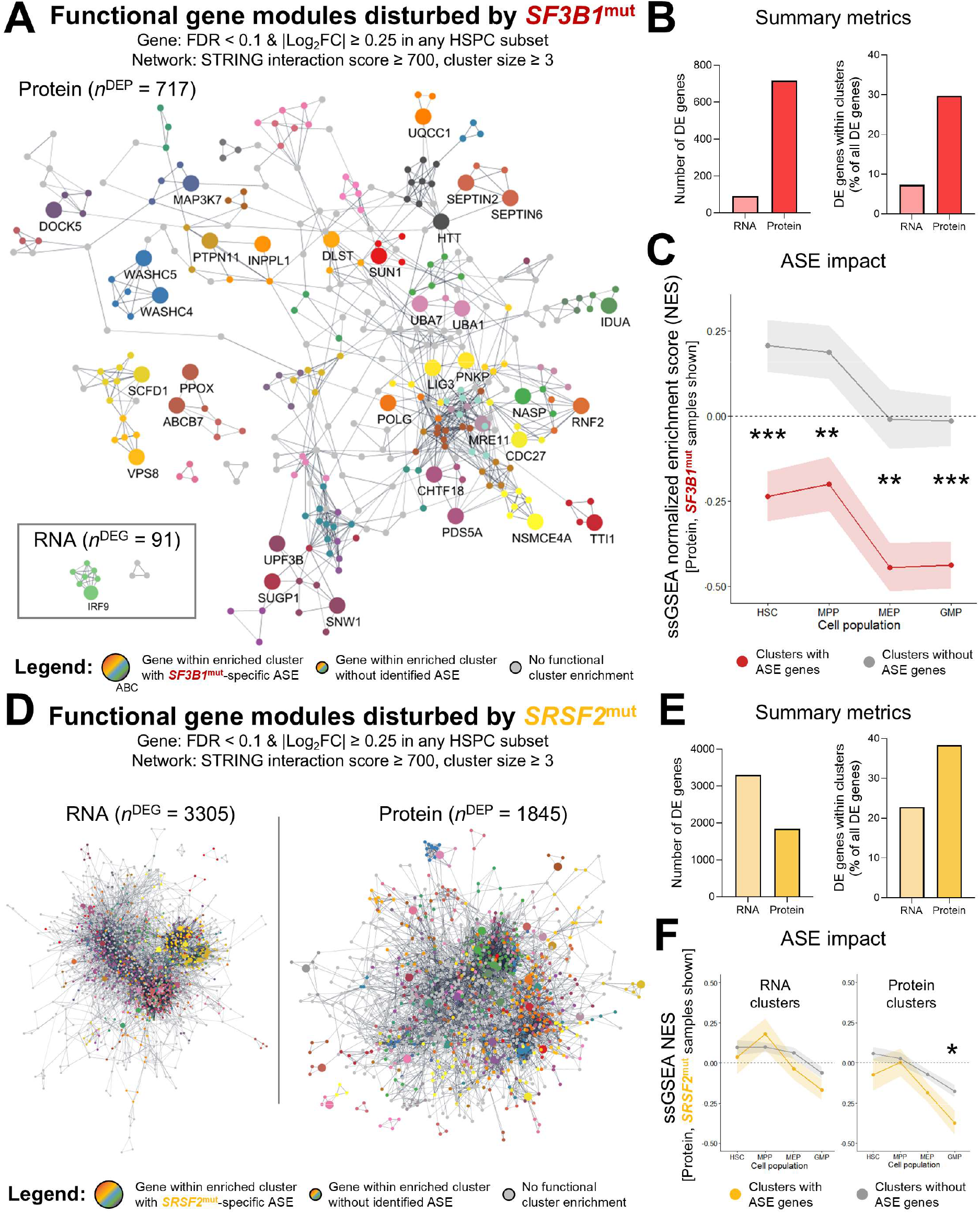
Aberrant RNA splicing disrupts protein interaction networks by protein complex poisoning without transcriptomic compensation. **A)** STRING protein-protein interaction networks (STRING interaction score ≥ 0.7) of *SF3B1*^mut^-associated differentially expressed genes (FDR < 0.10, |Log2FC| > 0.25) within the RNA layer (bottom-left box, n^DEG^ = 91) and the protein layer (overall landscape, nDEP = 717) as compared to a healthy donor background. Singleton nodes were removed from this visualization. Nodes with only one undirected edge were pruned from the network until reaching a minimum number of two undirected edges per node. Local network clusters were identified as fully connected subsets of proteins (maximal cliques) within the STRING interaction network. Cliques sharing proteins were iteratively merged until reaching a maximum Jaccard similarity index of 0.5 for each pair of cliques, and filtered to include only clusters with described protein complex identity as well as enrichment within the Gene Ontology database. **B)** Summary metrics of differentially expressed gene counts (left) and gene-cluster mapping percentages (right) within RNA and protein data of *SF3B1*^mut^ *vs.* healthy donor samples. **C)** Mean single sample gene set enrichment analysis (ssGSEA) normalized enrichment scores (NES) of all protein clusters within *SF3B1*^mut^ proteomic samples, comparing clusters containing genes with *SF3B1*^mut^-specific alternative splicing events (ASE genes) [red] against clusters without ASE genes (grey). Mean NES values were compared between ASE and non-ASE containing clusters through Student’s *t*-test. **D)** STRING protein-protein interaction networks of *SRSF2*^mut^-associated differentially expressed genes within the RNA layer (left, n^DEG^ = 3305) and the protein layer (right, nDEP = 1845) as compared to a healthy donor background. Network filtering was performed as described in A). **E)** Summary metrics of differentially expressed gene counts (left) and gene-cluster mapping percentages (right) within RNA and protein data of *SRSF2*^mut^ *vs.* healthy donor samples. **F)** Mean single sample gene set enrichment analysis (ssGSEA) normalized enrichment scores (NES) of all RNA clusters (left) and protein clusters (right) within *SRSF2*^mut^ proteomic samples, comparing clusters containing genes with *SRSF2*^mut^-specific alternative splicing events (ASE genes) [yellow] against clusters without ASE genes (grey). Mean NES values were compared between ASE and non-ASE containing clusters through Student’s *t*-test. * = p < 0.05, ** = p < 0.01, *** = p < 0.001.

An important internal validation of these data is the highly significant dysregulation of the UQCC1 chaperone trimer and mitochondrial complex III (**Fig. S7**), previously mechanistically associated with *SF3B1*^mut^ mis-splicing of *UQCC1.*^29^ However, whilst the UQCC1 trimer was underexpressed at protein level from HSC onward, mitochondrial complex III proteins were only significantly decreased in MEP and further dysregulated within the overall CD34^+^ HSPC compartment. This pattern of progressive dysregulation occurred in many of the identified protein complexes (*e.g*. Smc5-Smc6 complex, condensin complex, septin complex, TTT complex) and occurred primarily in association with RNA mis-splicing (**Fig. 3C**). Notably, few protein complexes were fully disrupted in HSC.

In comparison with the *SF3B1*^mut^ RNA network, *SRSF2*^mut^ were associated with much stronger patterns of differential RNA expression (*n*^DEG^ = 3305) (**Fig. 3D**). Despite this, similarly to *SF3B1*^mut^, *SRSF2*^mut^-associated differentially expressed proteins (*n*^DEP^ = 1845) displayed higher connectivity (**Fig. 3E**). Further, while clusters identified within RNA data failed to map to meaningful proteomic effects, *SRSF2*^mut^ protein clusters again displayed a pattern of progressive proteomic dysregulation, as described for *SF3B1*^mut^ (**Fig. 3F**).

Again, the reliability of this analysis is validated by the unbiased identification of the previously described integrator complex^23^ and EZH2/EED complex^27^ as disturbed functional clusters (**Fig. S8**). However, surprisingly, the overall levels of both integrator and EZH2/EED complexes were entirely stable in HSCs, with differentiation along the GMP trajectory (and CD34^+^ HSPC, primarily GMP in *SRSF2*^mut^ MN) being the main driver of protein complex disruption.

Thus, pauciproteomics reveals a novel pleiotropic layer of RNA mis-splicing effects, whereby ASEs directly impact the transcript-encoded protein in early cell stages, including HSCs, and gradually dysregulate (“poison”) functional protein complexes during differentiation.

### Proteotranscriptomic dynamics of the cancer-promoting SF^mut^ hematopoietic stem cell

The above identification of gradual proteome network effects justifies that direct analysis of the HSC compartment is the only means to identify putative mechanisms for SF^mut^ clonal expansion and MN development. This creates a challenge for functional validation, since modulation of true human HSC is not feasible, and mouse models do not recapitulate the RNA mis-splicing defects of human SF^mut^ (due to evolutionary intronic variation).

Thus, we pursued functional validation of early hematopoietic proteome phenotypes in induced pluripotent stem cell (iPSC) models to determine whether our MN observations remain valid in isogenic SF^mut^/SF^wt^ settings (**Data S7**). This approach focused on *SF3B1*^mut^ and *SRSF2*^mut^, where our proteotranscriptomic data had greater statistical power. For *SF3B1*^mut^ modelling, already known to be a cancer driving-mutation in isolation within the HSC compartment,^8^ we utilized a previously published and already characterized set of iPSC lines from our constellation, originally generated from a female MDS patient.^17^

Defining the role of *SRSF2*^mut^ in cancer promotion, on the other hand, poses a much greater challenge, since it is nearly always co-dominant with other driver mutations in our study cohort (**Fig. 1C**) and literature.^21^ We aimed to precisely define the specific effects imparted by *SRSF2*^mut^ alone within the cell; thus, in contrast with the previous strategy, we generated heterozygous *SRSF2*^P95H^ iPSC lines from a male healthy donor through CRISPR/Cas9 knock-in by homology directed repair. Both sets of isogenic lines were differentiated using a recently published definitive HSC-like iPSC differentiation protocol^30^ and later subjected to FACS for purification of CD34^+^CD38^-^early multipotent cells, obtaining a cell population representative of a multipotent HSC/MPP phenotype (**Fig. 4A**).

**Figure 4:**
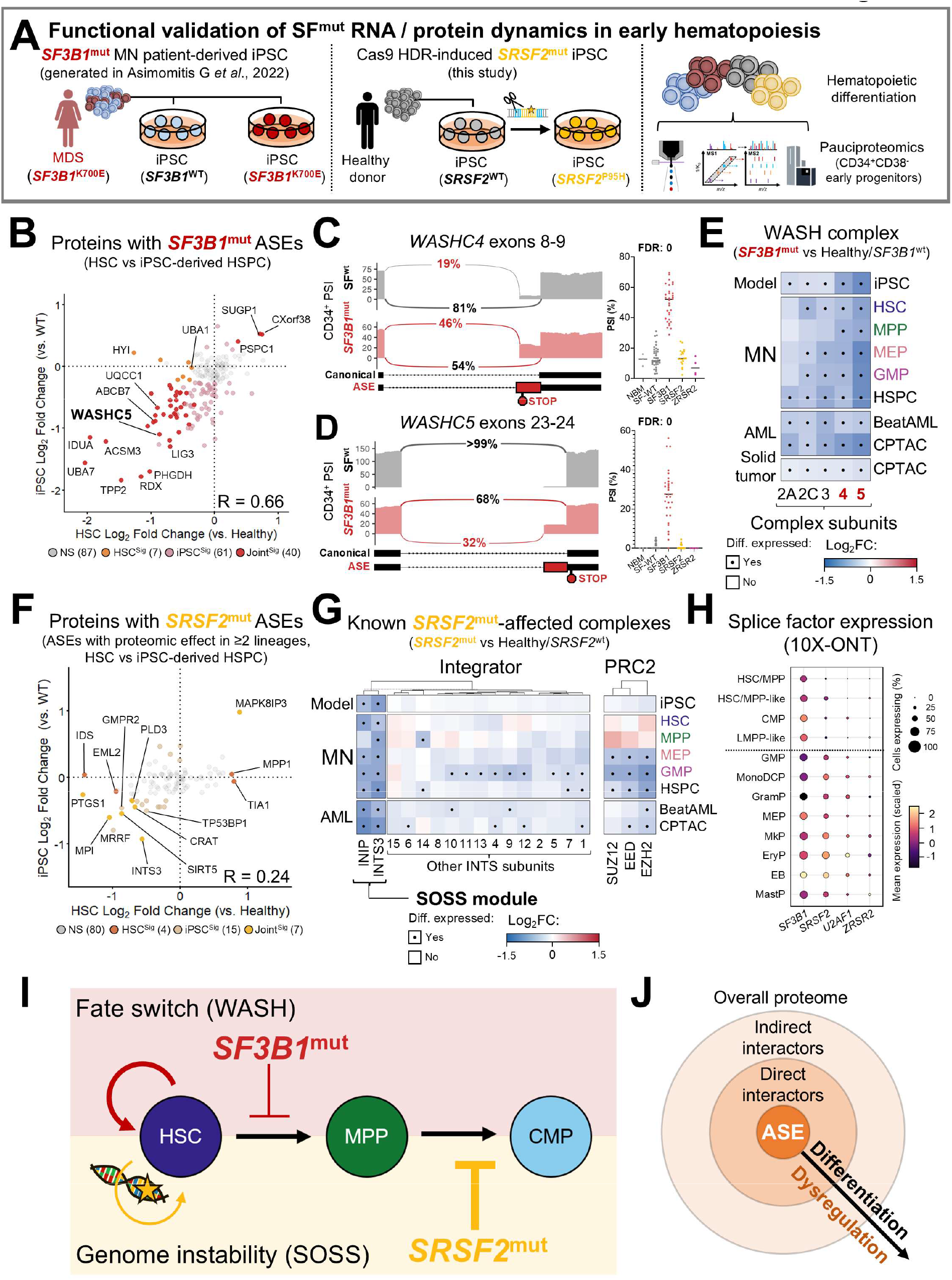
Induced pluripotent stem cell model-derived multipotent progenitors delineate mechanisms for the *SF3B1*^mut^ and *SRSF2*^mut^ stem cell advantage. **A)** Design of induced pluripotent stem cell (iPSC) culture experiments for functional validation of early hematopoietic proteome dynamics in the *SF3B1*^mut^ and *SRSF2*^mut^ settings. FACS-purified iPSC progenitors were defined by CD45^+^, CD34^+^ and CD38^-^ surface immunophenotypes (iPSC-derived MPP). **B)** Scatter plot of ASE-associated proteomic effects in *SF3B1*^mut^ HSC (X-axis) and *SF3B1*^mut^ iPSC-derived MPP (Y-axis). Significance cut-offs FDR < 0.10, |Log2FC| > 0.25. **C/D)** Sashimi plots of canonical (black) and ASE (red) PSI values for significant *SF3B1*^mut^ ASEs affecting *WASHC4* (**C**) and *WASHC5* (**D**). PSI values from bulk transcriptomic validation are shown to the right of each sashimi plot. **E)** Relative expression of WASH complex subunits within iPSC-derived MPP (top), MN HSPC subsets (middle) and external *SF3B1*^mut^ data (bottom, in order from the BeatAML 1.0, CPTAC-AML, and CPTAC solid tumor cohorts). Columns correspond to individual genes, rows correspond to cell compartments analyzed through pauciproteomics of iPSC models (iPSC), pauciproteomics of SF^mut^ MN HSPC (MN), tandem mass tag-based proteomics of acute myeloid leukemia from the BeatAML and CPTAC cohorts (AML), or tandem mass tag-based proteomics of solid tumors from the pan-cancer CPTAC cohort (Solid tumor). Log_2_ fold change (Log_2_FC) values are colored from blue (negative Log_2_FC, lower expression in mutant cells) to red (positive Log_2_FC, higher expression in mutant cells). Dots indicate differential expression significance in a given cell compartment or dataset (Pauciproteomics: FDR < 0.10, |Log2FC| > 0.25; TMT proteomic datasets: *p* < 0.10, |Log2FC| > 0.25). **F)** Scatter plot of ASE-associated proteomic effects in *SRSF2*^mut^ HSC (X-axis) and *SRSF2*^mut^ iPSC-derived MPP (Y-axis). **G)** Relative expression of Integrator (left) and Polycomb Repressive Complex 2 (PRC2, right) protein complex subunits within iPSC-derived MPP (top), MN HSPC subsets (middle) and external *SRSF2*^mut^ data (bottom, in order from the BeatAML 1.0 and CPTAC-AML cohorts). **H)** Dot plot of scaled RNA splice factor expression in 10X-ONT data grouped by labelled cell compartment. **I)** Diagram describing candidate advantage mechanisms in *SF3B1*^mut^ HSC (WASH complex loss and resulting fate conversion to further HSC accumulation) and *SRSF2*^mut^ HSC (increased genomic instability and significant blockade of downstream differentiation). **J)** General mechanism for SF^mut^ ASE-induced effects on the proteome.

The significant HSC proteomic dysregulation induced by *SF3B1*^mut^ RNA mis-splicing was generally well recapitulated in iPSC-MPP (**Fig. 4B**). Compared to the broader proteome network effects of later differentiation, *SF3B1*^mut^ impact only three functional protein complexes in HSC, all of which were recapitulated in iPSC-MPP (**Fig. S9A**). These were the UQCC chaperone trimer,^29^ albeit with normal levels of mitochondrial complex III at this stage; the single-strand break repair (SSBR) complex, wherein mis-splicing and protein loss of *LIG3* and *PNKP* disturbed the level of the critical DNA repair protein XRCC1; and the Wiskott Aldrich Syndrome protein and scar homologue complex (WASH complex). Specifically, RNA mis-splicing of *WASHC4* and *WASHC5*, previously not reported in detail, was highly specific to the *SF3B1*^mut^ setting and led to loss of the entire protein complex in HSC, iPSC-MPP and every external dataset evaluated, including pan-cancer solid tumors with *SF3B1* hotspot mutations (*WASHC4/5* mis-splicing events visualized in **Fig. 4C/D**, protein data in **Fig. 4E**).

Establishing these protein complexes as disturbed in HSC enables mechanistic associations with clonal expansion. Specifically, SSBR complex and XRCC1 dysregulation within HSC would be a potential cause for aberrant DNA repair in the *SF3B1*^mut^ setting, including genomic and telomeric instability with long-term consequences for clonal selection in aged hematopoiesis.^31,32^ The second association, which is especially relevant to the long-term clonal expansion of *SF3B1*^mut^, is that WASH complex deficiency has been mechanistically defined to impair the differentiation of long term-repopulating HSCs by dysregulating *c-Myc* activation, creating a strong competitive advantage in transplantation.^33^

Next, we analyzed *SRSF2*^mut^ iPSC-MPP. Similarly to primary MN HSC/MPP, iPSC-MPP displayed a mildly affected proteomic phenotype, with a nearly identical small set of ASE-associated proteomic events and in stark contrast with the defects observed at the GMP stage (**Fig. 4F**). Indeed, analysis of the total HSC proteome identified only one disturbed complex: the SOSS complex, a small multiprotein complex which includes the mis-spliced gene *INTS3*^23^ and its interactor *INIP*, creating a distinct module from the integrator/INTAC complex (**Fig. S9B**).^34^ As expected from the overall protein network analysis, integrator complex and EZH2/EED complex effects were absent in both HSC and iPSC-MPP (**Fig. 4G**). On the other hand, many of the dysregulated proteins in HSC/MPP and iPSC-MPP were exclusive to these earlier stages of differentiation (**Fig. S9C**), including deficiency of the critical hematopoietic transcription factor NFATC2.^35^ Importantly, loss of the SOSS complex alone can disable recruitment of the integrator complex,^34^ potentially triggering genomic instability in HSC.

However, the weak effect of *SRSF2*^mut^ in isolation led us to hypothesize that *SRSF2* could be differentially regulated during hematopoiesis. Indeed, while *SF3B1* was universally expressed at similar levels, *SRSF2* displayed significantly higher RNA as well as protein expression in later lineage-committed hematopoietic differentiation stages (**Fig. 4H**). Thus, in contrast with the ability of *SF3B1*^mut^ to drive disease in isolation, these proteotranscriptomic data justify the limited role of *SRSF2*^mut^ as isolated HSC driver mutations and further defines their role in the dysregulation of later hematopoietic stages, including AML transformation.

In summary, RNA mis-splicing caused by both *SF3B1*^mut^ and *SRSF2*^mut^ introduces separate dynamic proteotranscriptomic aberrations in HSC which can be mechanistically linked to disease-driving characteristics and cell phenotypes of these mutations, providing candidate explanations for their clonal expansion and progression mechanisms (**Fig. 4I**). Further, the combination of 10X-ONT and pauciproteomics establishes a causal sequence for how RNA mis-splicing ASEs gradually poison the proteome, leading to functional dysregulation (**Fig. 4J**).

## Discussion

Despite decades of transcriptomic studies, the cancer-initiating mechanisms of RNA splicing factor mutations have remained elusive. The clonal expansion and takeover enacted by SF^mut^ HSCs within the *in vivo* bone marrow portrays a striking contrast to the weak competitive ability of SF^mut^ cells in any model system or culture experiment. Further, the absence of steady-state human HSC model systems precludes the study of how individual mis-splicing events or protein defects contribute to the long-term expansion of human SF^mut^ HSCs.

Through integration of state-of-the-art single-cell RNA sequencing strategies, high-throughput low-cell proteomics and induced pluripotent stem cell modeling, the resulting data can now provide an explanation to this paradox. Specifically, SF^mut^ directly dysregulate individual proteins within the HSC through mis-splicing of the corresponding RNA transcripts, but do not substantially hamper the overall protein network at this stage. In contrast, upon early hematopoietic differentiation, these individual protein deficiencies progressively disturb numerous protein complexes and poison the overall protein network.

Notably, RNA mis-splicing disturbs certain protein complexes already within the HSC compartment with consequences for the clonal dynamics of myeloid neoplasms. We find that *SF3B1*^mut^ dysregulate WASH and SSBR complexes, connected to HSC fate determination^33^ and telomeric instability, respectively,^31,32^ justifying the role of *SF3B1*^mut^ in long-term oncogenesis as a sole driver mutation. *SRSF2*^mut^, on the other hand, dysregulate the SOSS module irrespectively of the integrator complex, which would trigger genomic instability in HSC and explain the accumulation of co-mutations.^34^ While this study could not yet determine cancer-driving mechanisms of *U2AF1*^mut^ and *ZRSR2*^mut^, we expect that future studies focusing on larger cohorts of these rare cases will be able to address these questions.

The gradual poisoning of the overall protein network may explain why SF^mut^ drive slow, long-term clonal expansion in HSC-driven myeloid malignancies and clonal hematopoiesis,^36^ in contrast with their catastrophic risk profile as a contributing mutation.^23,37^ Acute myeloid leukemia, chronic lymphocytic leukemia and solid tumors are all situations where cells without stem cell identity may regain self-renewal. SF^mut^ may drive much wider disruption of the proteome network in these diseases, which could accelerate clonal selection processes. Future SF^mut^ studies in different cancer settings, including a more detailed understanding of the order and effect of co-mutations, will be critical to precisely establish how SF^mut^ co-mutant tumorigenesis differs from long-term clonal HSC expansion.

This study establishes pauciproteomics as a high-throughput analytical method. Compared to the current state-of-the-art, pauciproteomics enables a large increase in sample throughput and proteome depth whilst maintaining the resolution needed to study minute cell populations. Thus, we provide a scaffold for future studies of cell differentiation, cancer, genetic model systems, and any context where the analysis of rare cell populations may be needed (or preferred, e.g. for high-throughput screening approaches). Indeed, pauciproteomics resolved healthy and SF^mut^ hematopoiesis in-depth, functionally defining how SF^mut^ ASEs impair the proteome at yet unachieved scale, and bypassing the need for cumbersome and cell number-heavy methods.

In conclusion, this study builds a generalizable proteotranscriptomic workflow for deep molecular investigation of rare cell populations, through which we have defined important pleiotropic effects of RNA splicing factor mutations in the hematopoietic system and elucidate their role in oncogenesis and disease.

## Methods

### Study design and ethical approval

Viably frozen bone marrow (BM) mononuclear cell (MNC) samples from SF^mut^ MN patients and healthy donors were requested and kindly received from the Karolinska Institutet Biobank, Stockholm, Sweden (*n* = 19 MN patients, n = 15 healthy donors [1 donor with 2 visits]); the Finnish Hematology Registry and Clinical Biobank, Helsinki, Finland (*n* = 29 MN patients); and the Kyoto University Biobank (*n* = 14 MN patients). As part of local clinical routines, MN patients from each site were genetically investigated with Next-Generation Sequencing panels for targeted sequencing of known myeloid malignancy driver genes (KI: GMS-Myeloid, 195 genes; FHRB: HUS-MyelMut, 68 genes; KU: In-house panel using the SureSelect custom kit [Agilent], 331 genes). Genes included in at least two NGS panels are listed in **Table S1**. The criteria for MN patient inclusion were: 1) a previous diagnosis of myelodysplastic syndromes (MDS), acute myeloid leukemia with MDS-related changes (MDS-AML) or chronic myelomonocytic leukemia (CMML), in accordance with local diagnostic criteria; 2) NGS-based genetic evaluation performed at diagnosis and/or follow-up, identifying at least one SF^mut^ with variant allele frequency ≥ 15%; 3) complete data concerning hematological parameters and survival outcomes; and 4) absence of multiple independent SF^mut^ clones. All source material was provided with written informed consent for research use, given in accordance with the Declaration of Helsinki, and the study was approved by the Ethics Research Committee at Karolinska Institutet (2017/1090-31/4, 2022-03406-02, 2024-03119-02). A deidentified donor index is provided in **Data S1**.

### Illumina/Nanopore integrated 10X Chromium single-cell RNA sequencing

Cryopreserved MNC were thawed, prepared for fluorescent labelling, analyzed and purified through fluorescence-activated cell sorting (FACS) using a FACS ARIA II Fusion (Becton Dickinson) at the MedH FACS facility in Karolinska Institutet. Antibodies used for sample labelling are listed in **Table S2**. Lin12^-^CD34^+^ live cells were viably purified onto tubes containing 10X Genomics resuspension buffer (PBS + 0.04% bovine serum albumin [Sigma]), left as single donor tubes or mixed at 50/50 male/female donor cell ratio for label-free sample multiplexing (**Figure S2A, Data S1**), and resuspended to a final concentration of 1,500 cells/µL, targeting a recovery of 15,000 cells/sample with the Chromium GEM-X Single-cell 3’ v4 kit (10X Genomics). The Chromium kit protocol was followed with modifications to increase cDNA length. Libraries were pooled and sequenced on an Illumina NovaSeq 25B (Illumina), read length 100 bp, or prepared individually for sequencing on a PromethION system (Oxford Nanopore Technologies), protocol version SST_9198_v114_revO, achieving a mean read length of ∼900 bp across all samples. Extended details are provided in **Sup. Methods**.

### Low-cell proteomics sample processing

Cryopreserved MNC were prepared as above and FACS-purified onto the center of dry Protein LoBind tubes (Eppendorf) using a FACS ARIA II Fusion (Becton Dickinson) at the MedH FACS facility of Karolinska Institutet (sorting strategy in **Fig. S3A**). After sorting of up to a maximum of 1000 cells, samples were snap frozen in dry ice and later prepared for mass spectrometry analysis using a procedure of digestion on Evotip based on and modified from Ye et al.^38^. Finally, the samples were analyzed for LC-MS/MS using a Evosep One (Evosep) coupled to a timsTOF SCP mass spectrometer (Bruker) in data independent acquisition diaPASEF (parallel accumulation-serial fragmentation) mode. Extended details are provided in **Sup. Methods**.

### Data analysis and statistical methods

Single-cell RNA sequencing: Illumina data files were processed using CellRanger v.9.0.1 (10X Genomics). Nanopore data files were processed with the EPI2ME wf-single-cell workflow v.3.3.0 (Oxford Nanopore Technologies). Both sets of files were aligned against the GRCh38 reference genome. Raw mass spectrometry data files were analyzed using Spectronaut v.19.4 (Biognosys) with the DirectDIA workflow for label-free quantification. Extended details are provided in **Sup. Methods**. This study also makes use of bulk RNA sequencing data previously reported in Shiozawa *et al*. 2018^22^ and generated by the Department of Pathology and Tumor Biology, Kyoto University. External cohort proteomics data used in this publication were generated by the National Cancer Institute Clinical Proteomic Tumor Analysis Consortium (CPTAC). Gene set enrichment analyses were performed via GSEA (v. 4.4.0)^39,40^ or the STRING database (v. 12.0).^28^ All statistical analyses were performed with RStudio v.1.4.1767, R v.4.0.5 and GraphPad Prism v.9.4.0.

### Induced pluripotent stem cell (iPSC) protocols

*SF3B1*^mut^ iPSC source: iPSC lines were established in Asimomitis *et al*.^17^ Isogenic *SF3B1*^WT^ and *SF3B1*^K700E^ lines correspond to the female MDS patient (P22) within the cited study.

*SRSF2*^mut^ line generation: The human 1157 iPSC line was transfected using the Lonza 4D cell line kit with ribonucleoproteins (RNPs) containing Cas9 and a guide RNA (gRNA) targeting the human *SRSF2* gene at residues encoding P95, as well as a single stranded homology directed repair template bearing the P95H mutation. After 72 hours, transfected cells were plated at single cell density and single cell clones were isolated. Clones were screened for introduction of heterozygous *SRSF2*-P95H mutation using Sanger sequencing. Heterozygous targeted and wild-type untargeted clones were obtained, pluripotency confirmed by immunostaining for pluripotency markers, and copy number variations screened using G-band karyotyping.

Culture: iPSC-derived HSCs were generated using a swirling embryoid body (EB) differentiation protocol.^30^ Briefly, iPSCs were dissociated into single cells using EZ-LiFT (Sigma-Aldrich), and 2×10^6^ cells were seeded in day 0 (D0) medium as described in protocol 3 of Ng *et al*.^30^ Medium was changed every other day, and floating HSCs were harvested on D14 by collecting the supernatant from the swirling EB cultures. CD34^+^CD38^-^ cells were purified for later mass spectrometry as described above.

## Acknowledgments

The authors would like to thank the MN patients and healthy donors for their willingness to participate in this research, as well as the crucial biobanking activities and sample access provided by the Karolinska Institutet Biobank (Stockholm, Sweden), the Finnish Hematology Registry and Clinical Biobank (Helsinki, Finland), and the Kyoto University Biobank (Kyoto, Japan). Additionally, the authors would like to acknowledge the Karolinska Institutet Bioinformatics Expression and Analysis Facility for assistance with Nanopore and Illumina sequencing, the Single-Cell Core Facility (SICOF) for assistance with the Chromium GEM-X protocol, and the MedH Flow Cytometry Core Facility for providing instruments for cell sorting and analysis. These facilities are financed by the Infrastructure Board at Karolinska Institutet. Mass spectrometry analyses were performed using instrumentation located at the Clinical Proteomics Mass Spectrometry facility, Karolinska Institutet / Karolinska University Hospital / Science for Life Laboratory. The authors acknowledge support from the National Genomics Infrastructure in Stockholm funded by Science for Life Laboratory, the Knut and Alice Wallenberg Foundation and the Swedish Research Council, and SNIC/NAISS/Uppsala Multidisciplinary Center for Advanced Computational Science for assistance with massively parallel sequencing and access to the UPPMAX computational infrastructure. PLM was supported by the Myelodysplastic Syndromes Foundation, Inc. (grant number 1142079), the European Hematology Association (Advanced Research Grant Topics-in-Focus Precision Hematology), the Dr. Åke Olsson foundation (Dnr 2024-00303), the Alex and Eva Wallström foundation (Dnr 2024-00311), the Jeanssons Stiftelse (Dnr 4-3521/2025), the Felix Mindus contribution to Leukemia Research (Dnr 2025-02863) and a KI Research Foundation grant (Dnr 2024-02330). CMP was supported by a Karolinska Institutet Doctoral funding grant (Dnr 2025-01553). VL was supported by Vetenskapsrådet (grant number 2020–01902), Cancer Research KI (Karolinska Institutet), and Felix Mindus contribution to Leukemia Research. EHL was supported by Cancerfonden (grant number 19 0200), Vetenskapsrådet (grant number 211133), Knut and Alice Wallenberg Foundation (grant number 2017.0359). RMB, YK and JL were supported by Vetenskapsrådet (grant number 2021-00754), Erling Persson Stiftelse (grant number 22/9-2020), Cancerfonden (grant numbers 20 1269 Pj F, 23 2819 Pj 01 H), VINNOVA (2024-01137), the Cancer Research Foundations of Radiumhemmet (grant number 241272), Knut and Alice Wallenberg Foundation (grant number 2024.0032) and the Stockholm County Council (SLL) Region Stockholm (grant number FoUI-1000396).

## Authorship contributions

PLM, RMB, JL and EHL conceived and designed the study. PLM, RMB, RGR, VL, PSW, SEWJ, YN, SO, JL and EHL contributed to experiment design and result interpretation. PLM, SH, CMP, AA, SF, MMN, TM-B, SB, CK, TS, IB and A-CB conducted laboratory experiments. RMB, YK and KRS conducted mass spectrometry experiments. MMN, TM-B, MC, IB, A-CB, MHW, JU, YN, SO and EHL conducted or administered clinical biobanking activities. SB, CK, TS and RGR created and validated *SRSF2*^mut^ iPSC lines. SH and CMP conducted iPSC cultures under supervision of VL. PLM and IS conducted computational analyses. DB, SM, MS and FF-B contributed to 10X-ONT design and optimization steps. SEWJ, JU, YN, SO, JL and EHL provided computational and sample resources. PLM, RMB, JL and EHL created figures and wrote the manuscript. All authors read, edited, and approved the manuscript.

## Data availability

Preprocessed data are in the supplemental material. Deidentified raw data from 10X-ONT (FASTQs, count matrices, BAM files and analysis-ready Seurat v5 objects) have been deposited to the Swedish National Data Service (SND) data repository with restricted access, in accordance with the European Union General Data Protection Regulation, and are accessible upon reasonable request to SND (SND 2026-285). Pauciproteomics data (raw files, search files, search parameters and fasta file) have been deposited to the ProteomeXchange Consortium via the PRIDE partner repository with the dataset identifier PXD065285. Other raw data are available from the corresponding authors upon request.

## MAIN SUPPLEMENTARY DOCUMENT

### Supplemental data descriptions

**Data S1**: Index of patients and healthy donors analyzed in this study, including genomic information and quality control metrics for the 10X-ONT and pauciproteomics datasets.

**Data S2:** Pre-processed 10X-ONT and pauciproteomics data for the healthy donor and MN patient cohort. A lineage-aggregated count matrix is provided for 10X-ONT. VSN-normalized MS2 data are provided for pauciproteomics.

**Data S3:** Marker gene expression of annotated cell populations in 10X-ONT data.

**Data S4**: Differential expression analysis of protein and RNA content in hematopoietic stem and progenitor cell compartments, comparing each SF^mut^ MN group against healthy donors.

**Data S5**: List of mis-spliced transcripts. Events were validated for *SF3B1*^mut^, *SRSF2*^mut^ and *ZRSR2*^mut^, and further collated from literature for *SRSF2*^mut^ and *U2AF1*^mut^. *SF3B1*^mut^ events are detailed in regard to splice site usage, protein sequence consequences and nonsense-mediated decay prediction.

**Data S6**: STRING network analysis of differentially expressed genes and resulting interactive networks.

**Data S7**: VSN-normalized MS2 data and differential expression analysis of isogenic pairs of *SF3B1*^K700E-MDS^ vs. *SF3B1*^WT-MDS^ and *SRSF2*^P95H^ vs. *SRSF2*^WT^ induced pluripotent stem cell samples (CD34^+^38^-^).

### Supplemental figure legends

**Figure S1:**
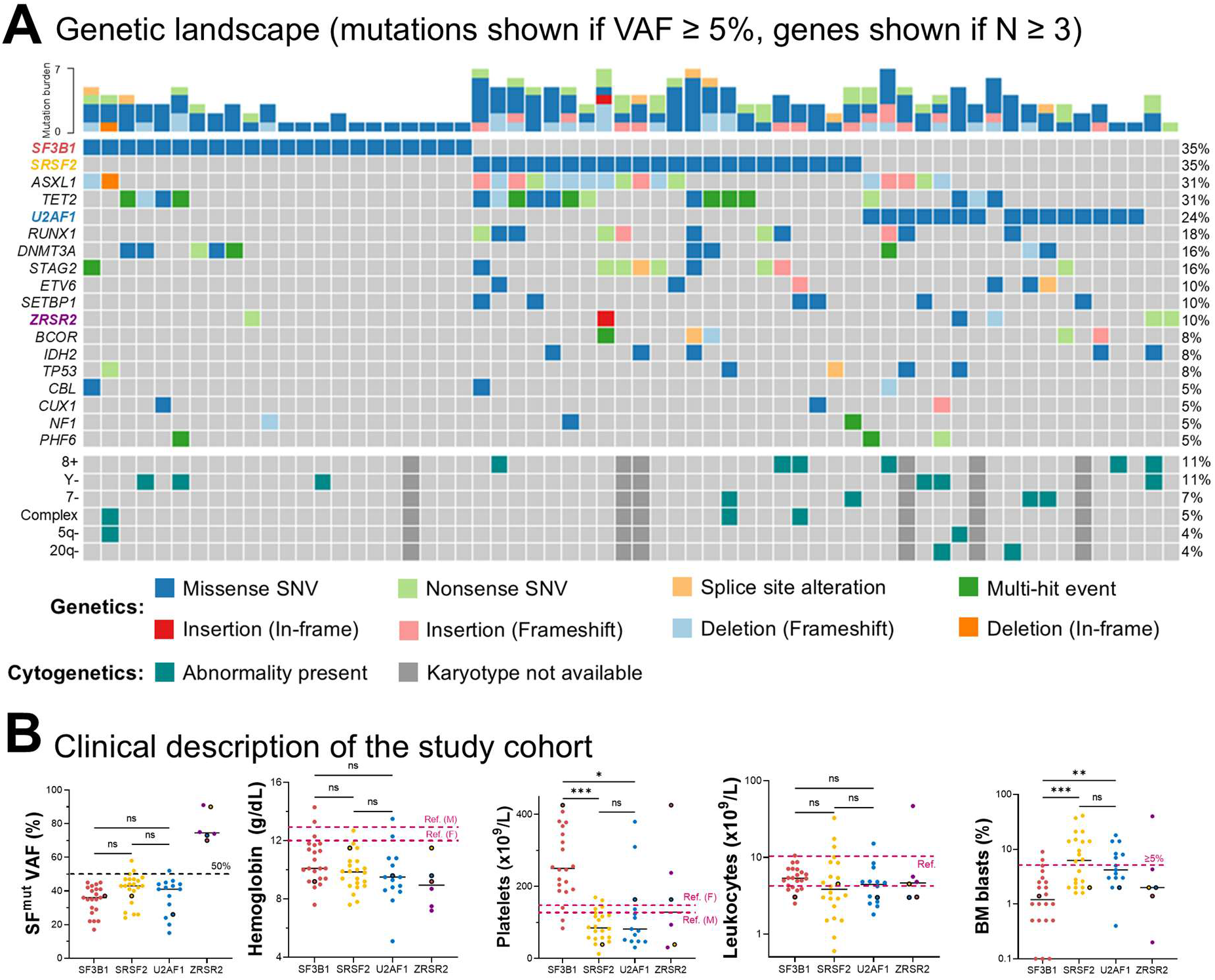
Clinical and genomic characterization of the SF^mut^ study cohort. **A)** Oncoplot showing mutations and cytogenetic abnormalities across the patient cohort, sorted by mutational frequency. The total number of mutations per patient is shown on the topmost bar plot. Only mutations occurring at VAF ≥ 5% and in at least three separate patients are shown. **B)** Dot plots showing clinical data of patients grouped by major SF^mut^ category (one dot per individual + median line) encompassing variant allele frequency as determined by diagnostic myeloid neoplasm panel sequencing of bone marrow mononuclear cells (VAF, dotted line at 50% indicates the break point for total heterozygosity), hemoglobin (male reference limit = 13.5 g/dL, female reference limit = 12.0 g/dL), platelet counts (male reference limit = 130 x 10^9^/L, female reference limit = 150 x 10^9^/L), leukocyte counts (min. reference limit = 4.5 x 10^9^/L, max. reference limit = 11 x 10^9^/L) and bone marrow blast percentages (5% break point for consideration as excess blast frequencies). *ZRSR2*^mut^ + SF^mut^ patient data is shown twice as black outlined circles colored according to the non-*ZRSR2* mutation (n = 1 per mutation, not considered for statistical comparisons). * = p < 0.05, ** = p < 0.01, *** = p < 0.001, ns = non statistically significant.

**Figure S2:**
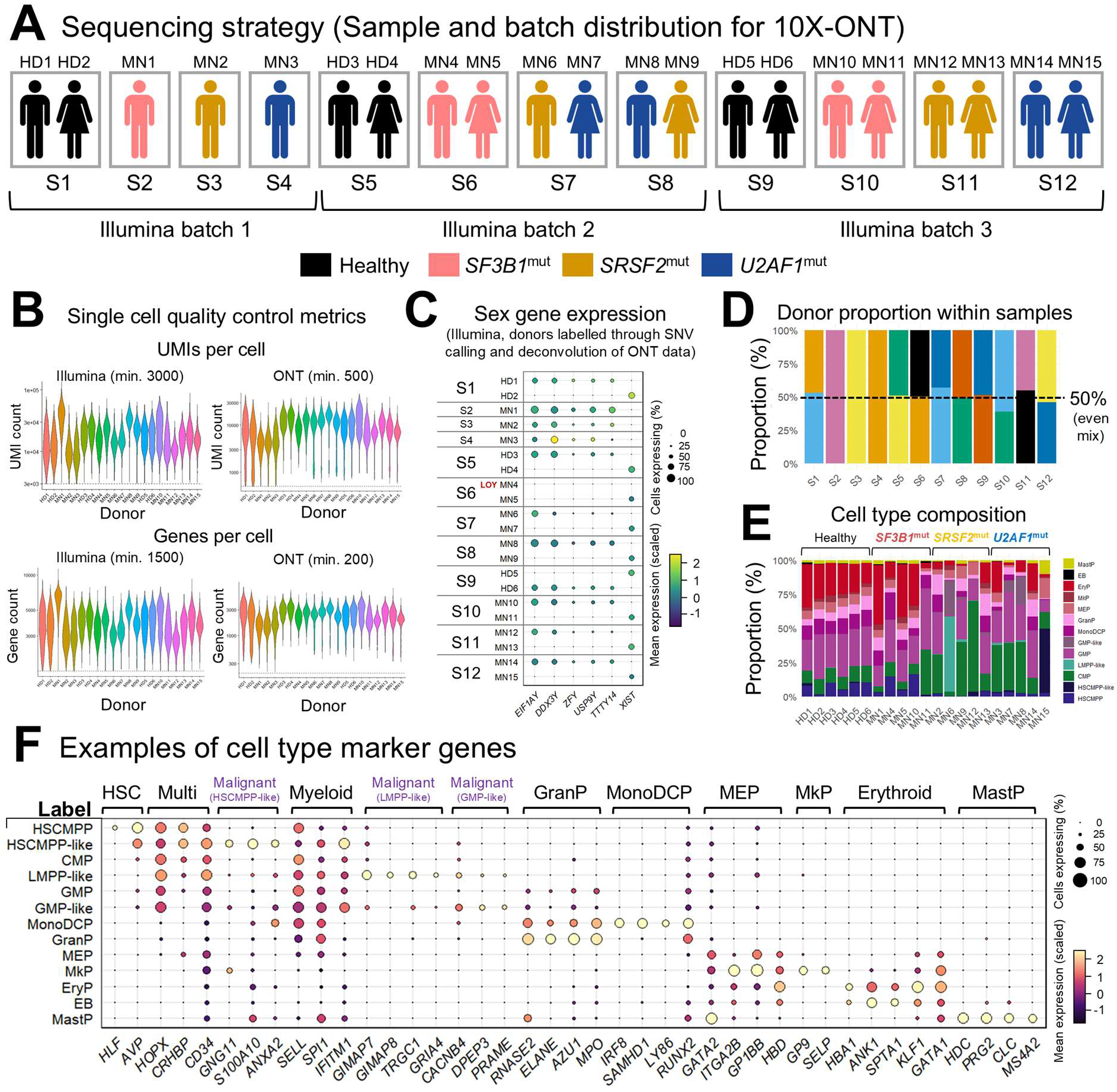
10X-ONT experiment design and quality control metrics. **A)** Sample distribution and design for 10X-ONT single-cell RNA sequencing (scRNAseq). Healthy donor (HD) and SF^mut^ MN patient (MN) bone marrow mononuclear cells were thawed and subjected to FACS for purification of Lin12^-^CD34^+^ live singlets. Where indicated, samples from donors of different sex were mixed at a 50/50 ratio before proceeding with the Chromium GEM-X 3’ v4 single-cell protocol in order to increase the number of biological replicates per group and enable downstream pseudobulk analyses. In total, 21 HD/MN biological samples (*n*^NBM^ = 6, *n^SF3B^*^1^ = 5, *n^SRSF^*^2^ = 5, *n^U2AF^*^1^ = 5) were processed as 12 Chromium samples. Each sample was sequenced in an individual PromethION flow cell as well as batched for a total of three Illumina sequencing rounds. Sample demultiplexing into individual donors was performed computationally through single nucleotide variation (SNV) calling with the Epi2me ONT analysis pipeline followed by clustering of donor-specific SNVs, and confirmed by analysis of sex-specific gene expression. **B)** Violin plots of obtained Unique Molecular Identifier (UMI) counts per cell and detected gene numbers per cell in Illumina-sequenced (left) and ONT PromethION-sequenced (right) 10X single-cell RNAseq data after quality control cut-offs were applied (Illumina: Min. 3000 UMI/cell and 1500 genes/cell, max 8% mitochondrial reads/cell. ONT: Min. 500 UMI/cell and 200 genes/cell, max. 20% mitochondrial reads/cell). **C)** Dot plot of sex gene expression across all SNV-demultiplexed donors. Male-specific genes: *EIF1AY*, *DDX3Y*, *ZFY*, *USP7Y*, *TTTY14* (Y chromosome-encoded genes). Female-specific gene: *XIST* (X-inactive specific transcript). Donor MN4 has lost the Y chromosome (LOY) as confirmed by clinical karyotyping (45,X-Y). **D)** Proportion of individual donor contribution to each sequenced sample after quality control and SNV-demultiplexing. **E)** Cell type composition per donor. **F)** Dot plot of marker gene expression for each labelled cell type. Labelled cell types are indicated on the left, marker gene descriptions are indicated on the top (e.g. *HLF* and *AVP* are HSC marker genes).

**Figure S3:**
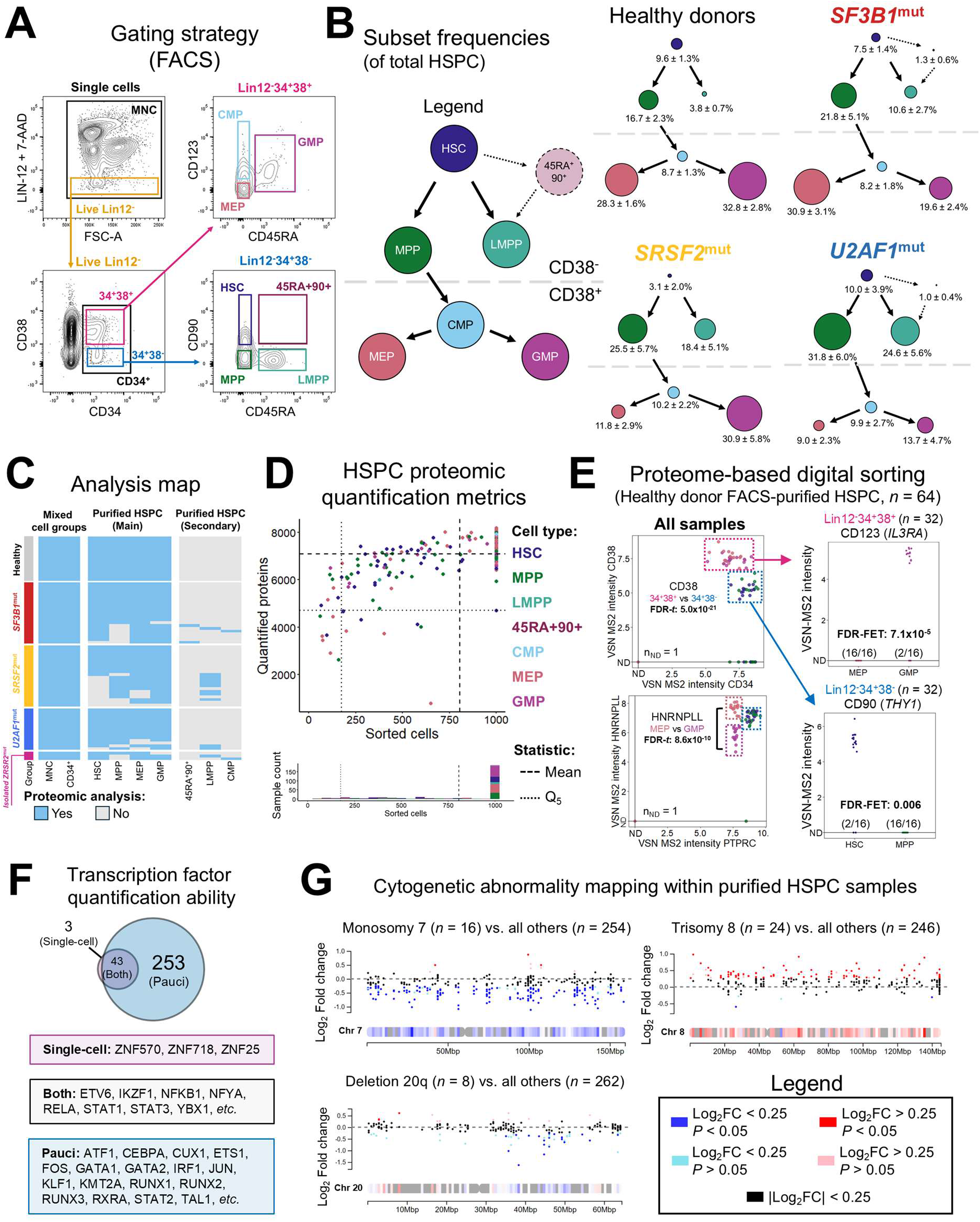
Pauciproteomics design and data quality assessment. **A)** Gating strategy for analysis, purification of hematopoietic stem and progenitor subsets and sample preparation. MNC = mononuclear cells; HSPC = hematopoietic stem and progenitor cells; HSC = hematopoietic stem cells; MPP = multipotent progenitors; LMPP = lymphoid-primed multipotent progenitors; 45RA+90+ = CD45RA and CD90 co-expressing CD38-negative cells; CMP = common myeloid progenitors; MEP = megakaryocyte-erythroid progenitors; GMP = granulocyte-monocyte progenitors. **B)** Summary of hematopoietic stem and progenitor cell frequencies along the hematopoietic hierarchy (Mean +/- SEM; *n^Healthy^* = 16, *n^SF3B^*^1^ = 22, *n^SRSF^*^2^ = 22, *n^U2AF^*^1^ = 15, *n^ZRSR^*^2^ = 6; patients with combined SF^mt^ + *ZRSR2*^mt^ were counted for both sets). The area of each quantified subset is scaled to its relative frequency within the sum of all phenotyped cells. **C)** Heatmap of samples processed for pauciproteomics. **D)** Quantified protein numbers in purified HSPC subsets of variable cell number, including sorted sample cell number distribution in the bar graph. Means indicated by dashed lines, 5^th^ percentiles indicated by dotted lines. **E)** Quantification of HSPC subset markers CD34, CD38, CD45 (PTPRC), HNRNPLL (splice factor controlling CD45RA expression), CD123 (IL3RA) and CD90 (THY1) in healthy bone marrow donor purified subsets. The number of samples where detection was not achieved (ND) is indicated at the bottom of each graph. **F)** Overlap of quantified transcription factors (as a known lower-abundance protein group) in single-cell proteomics (magenta, Furtwängler B *et al*., *Science* 2025) and pauciproteomics (blue). **G)** ChromoMap visualization of relative protein levels across the length of chromosomes 7 (top left), 8 (top right) and 20 (bottom left). The X-axis visualizes quantified proteins along each chromosome (distance annotated in million base pair counts [Mbp]). The Y-axis visualizes relative Log2 protein fold change between samples with cytogenetic abnormalities against all other samples.

**Figure S4:**
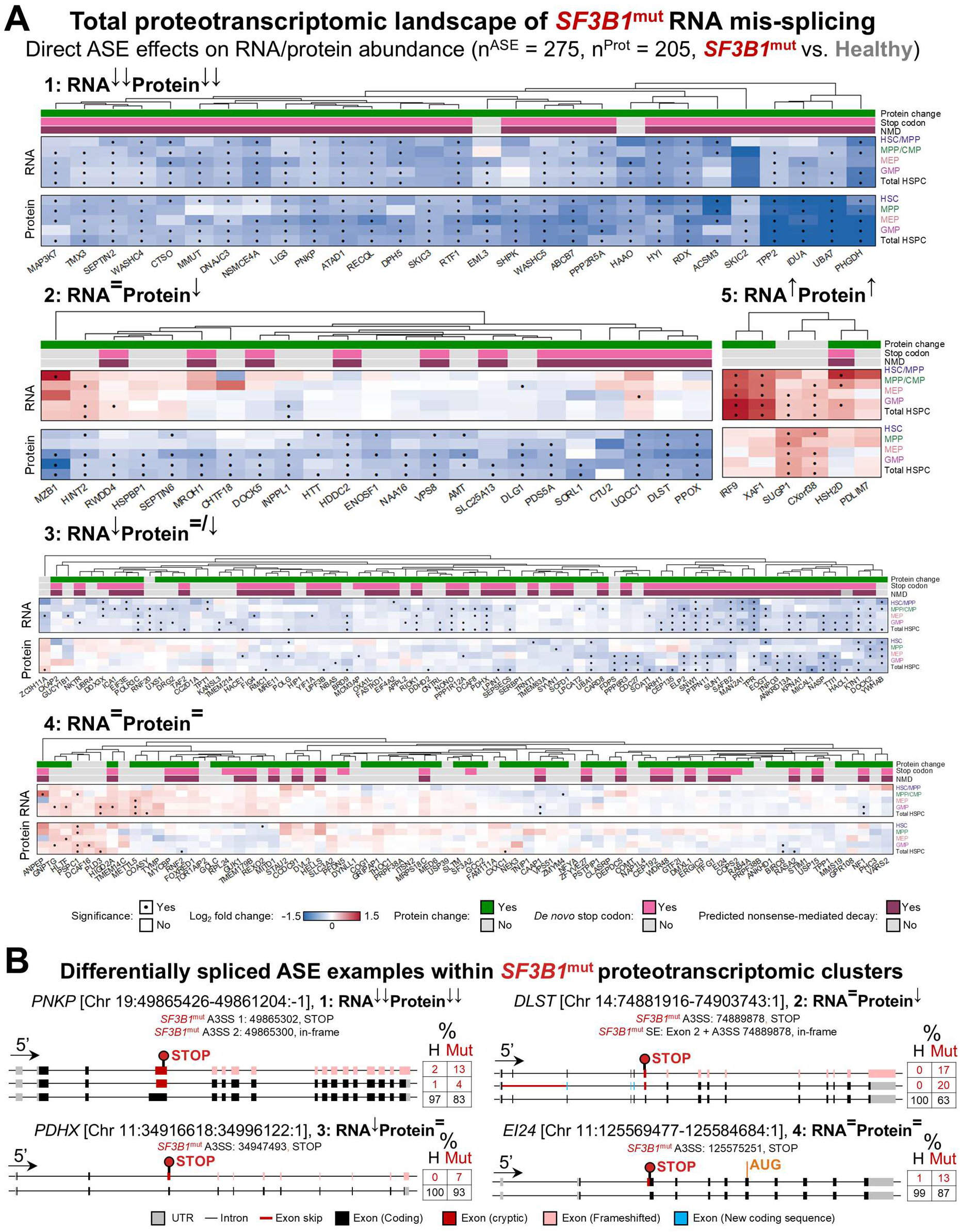
Detailed proteotranscriptomic landscape of *SF3B1^mut^* RNA mis-splicing. **A)** Relative RNA/protein heatmap (*SF3B1*^mut^ vs. Healthy) of genes with significant *SF3B1*^mut^ ASEs, separated by clusters of RNA/protein expression (obtained by unsupervised K-means clustering [K = 5]). Predicted ASE effects on the protein sequence, stop codon induction and nonsense-mediated decay (NMD) dynamics are marked above each heatmap. **B)** Examples of mis-spliced gene isoforms within major differential RNA/protein expression clusters, including genomic coordinates. Mean transcript isoform distribution is labelled to the right of each isoform in healthy (*H*) and *SF3B1*^mut^ samples (*Mut*).

**Figure S5:**
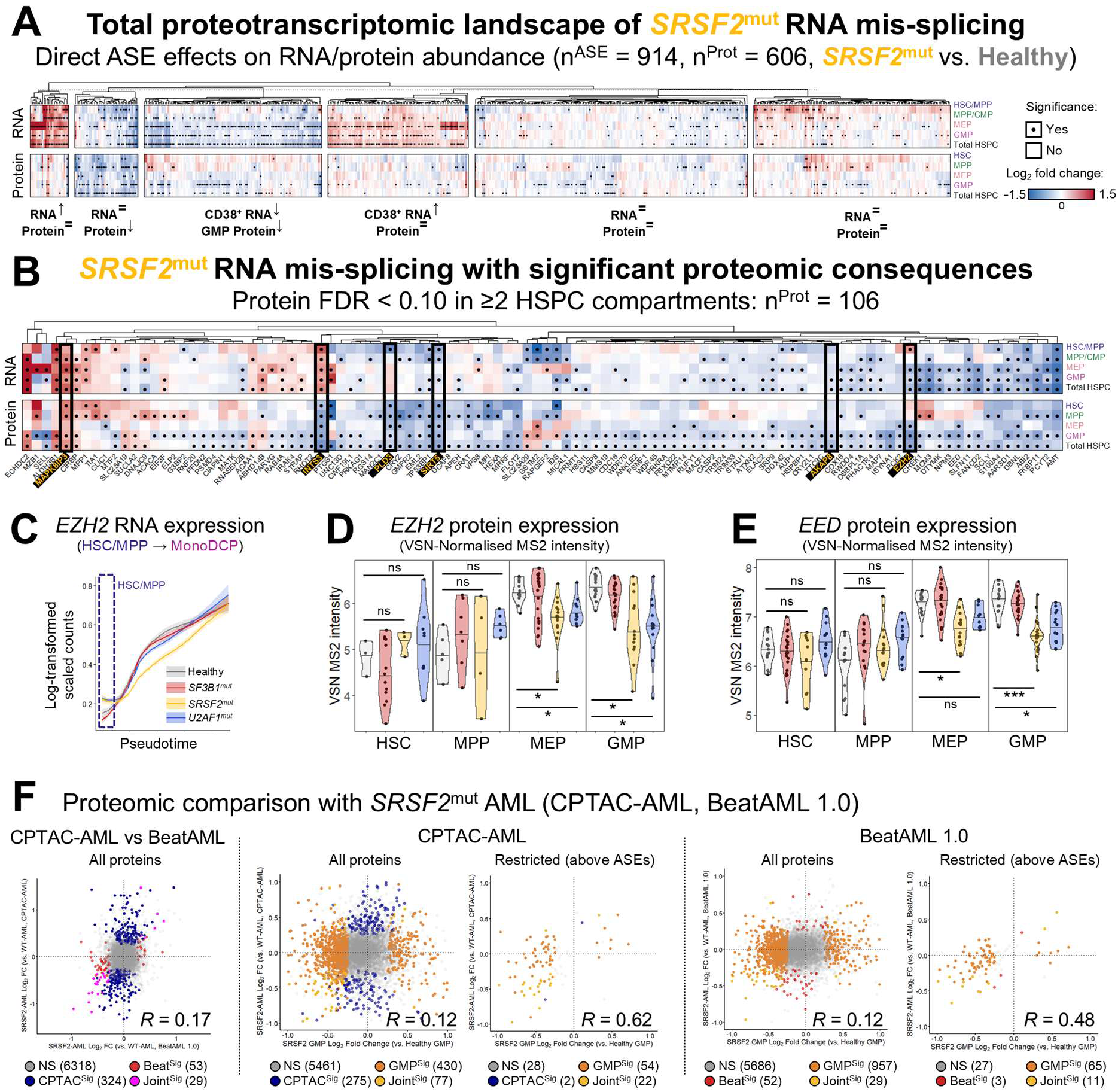
Detailed proteotranscriptomic landscape of *SRSF2*^mut^ RNA mis-splicing. **A)** Relative RNA/protein heatmap (*SRSF2*^mut^ vs. Healthy) of genes with significant *SRSF2*^mut^ ASEs, separated by clusters of RNA/protein expression (obtained by unsupervised K-means clustering [K = 6]). **B)** Relative RNA/protein heatmap of genes with significant *SRSF2*^mut^ ASEs which are associated with significant proteomic consequences within hematopoietic stem and progenitor cell compartments (FDR<0.10 in 2 or more compartments). Key genes affected by *SRSF2*^mut^ RNA mis-splicing are labelled with black and yellow text and highlighted with black boxes (from left to right: proteins affected in all stages: MAPK8IP3, INTS3, PLD3, SIRT5; proteins restricted to CD38^+^ cells: AKAP8, EZH2). **C)** Smoothed mean log-transformed scaled expression values of *EZH2* along pseudotime, separated by mutation. Expression curves were modelled using the Locally Estimated Scatterplot Smoothing (LOESS) method. **D/E)** Violin plots of **D)** EZH2 and **E)** EED protein expression. **F)** Scatter plots comparing MN data from this study against external AML cohorts which include *SRSF2*^mut^ cases. From left to right: 1) All proteins, CPTAC-AML (Y) vs. BeatAML (X); 2) All proteins, MN GMP (X) vs CPTAC-AML (Y). 3) Same comparison as 2) but with proteins restricted to the 106 proteins with SRSF2^mut^ ASE-associated proteomic effects; 4) MN GMP (X) vs BeatAML (Y); 5) Same as 4) but restricted to the same 106 proteins. Correlation coefficients (R) are indicated in each graph.

**Figure S6:**
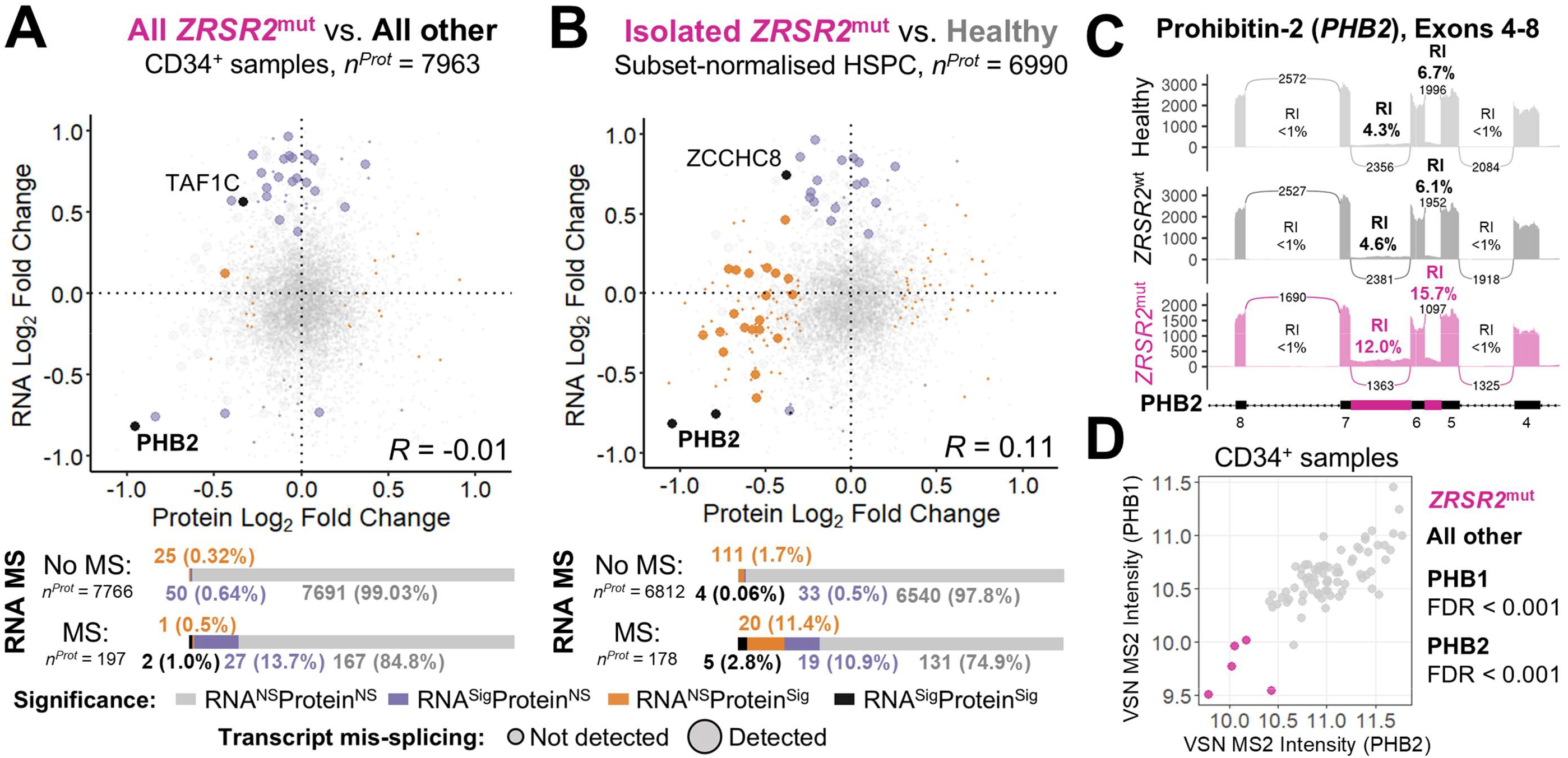
Proteomic analysis of *ZRSR2*^mut^ RNA mis-splicing effects. **A)** Correlation analysis of relative protein and RNA expression across the quantifiable proteome for *ZRSR2*^mut^ samples vs. all other samples in the CD34^+^ compartment and **B)** merged hematopoietic stem and progenitor cell subsets of *ZRSR2*^mut^ vs. healthy donor samples. The RNA data for this analysis was obtained from Shiozawa *et al*. 2018, comparing *ZRSR2*^mut^ MDS MACS-enriched CD34^+^ cells compared against SF^wt^ MACS-enriched CD34+ cells. **C)** Sashimi plot visualizing two *ZRSR2*^mut^-associated intron retention events in the PHB2 (Prohibitin-2) gene. **D)** Scatter plot of PHB2 protein expression vs. PHB1 (Prohibitin-1) protein expression in CD34^+^ HSPC samples. FDR-adjusted P-values are shown for each protein (*ZRSR2*^mut^ vs. all other).

**Figure S7:**
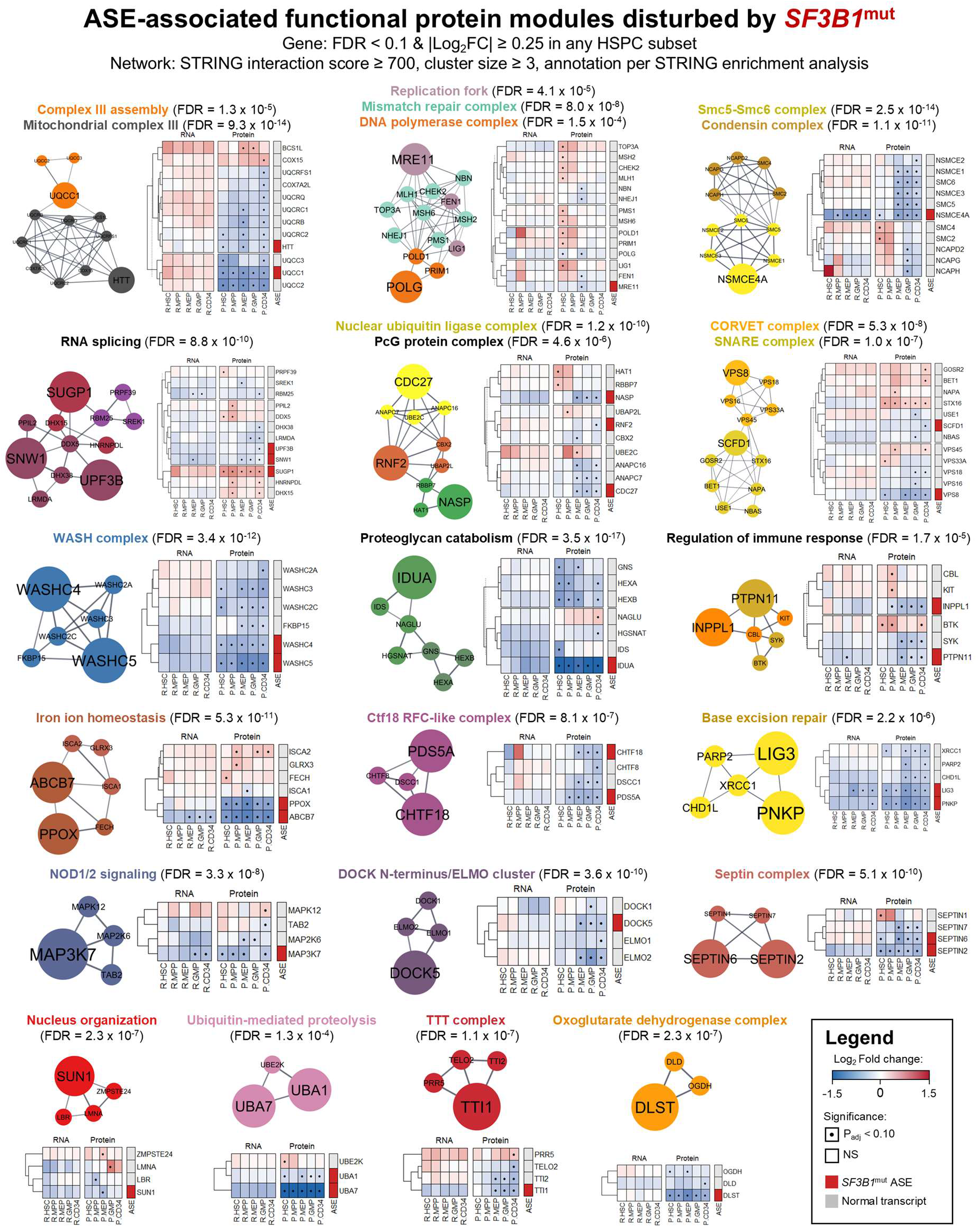
Mapping of differentially regulated protein complexes associated with *SF3B1*^mut^-specific RNA splicing. STRING protein-protein interaction networks of disturbed protein complexes within the *SF3B1*^mut^ setting and associated with *SF3B1*^mut^-specific alternative splicing events (ASE). Protein complex labelling was performed via gene set over-representation analysis using the STRING database. Protein complex identity associated with unique clusters is labelled with the respective cluster color. Functional enrichment data for the entire cluster is otherwise labelled in black. Relative RNA and protein heatmaps comparing *SF3B1*^mut^ against healthy donor samples are added to the right of each cluster. Within the heatmaps, Log2 fold change (Log2FC) values are colored from blue (negative Log2FC, lower expression in mutant cells) to red (positive Log2FC, higher expression in mutant cells). Dots indicate differential expression significance in a given cell compartment (FDR < 0.10, |Log2FC| ≥ 0.25). The boxes next to each gene indicate whether its transcript is modified by *SF3B1*^mut^-specific alternative splicing events (red = *SF3B1*^mut^ ASE, grey = normal transcript).

**Figure S8:**
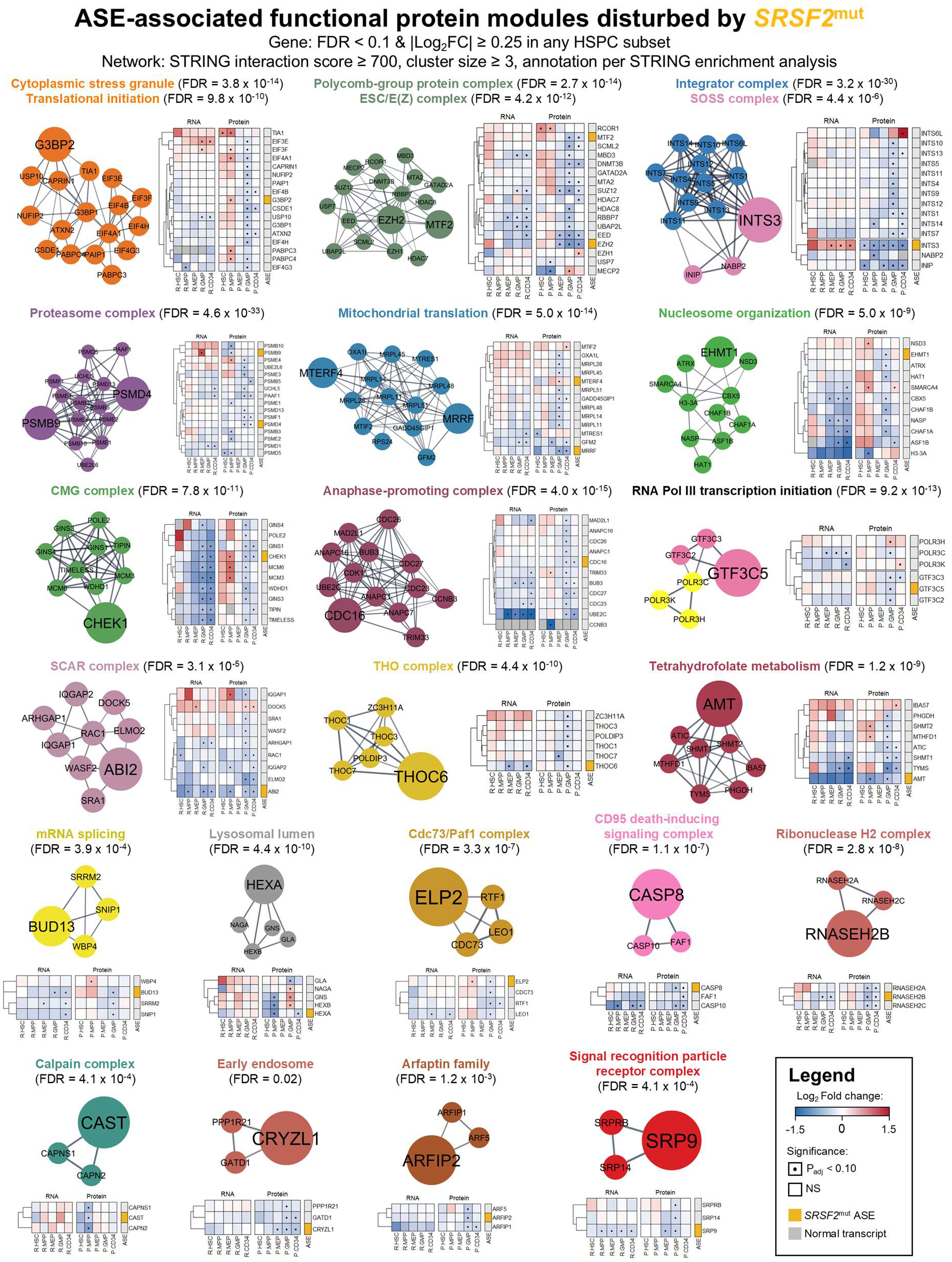
Mapping of differentially regulated protein complexes associated with *SRSF2*^mut^-specific RNA splicing. STRING protein-protein interaction networks of disturbed protein complexes within the *SRSF2*^mut^ setting and associated with *SRSF2*^mut^-specific alternative splicing events (ASE). Protein complex labelling was performed via gene set over-representation analysis using the STRING database. Protein complex identity associated with unique clusters is labelled with the respective cluster color. Functional enrichment data for the entire cluster is otherwise labelled in black. Relative RNA and protein heatmaps comparing *SRSF2*^mut^ against healthy donor samples are added to the right of each cluster. Within the heatmaps, Log2 fold change (Log2FC) values are colored from blue (negative Log2FC, lower expression in mutant cells) to red (positive Log2FC, higher expression in mutant cells). Dots indicate differential expression significance in each cell compartment (FDR < 0.10, |Log2FC| ≥ 0.25). The boxes next to each gene indicate whether its transcript is modified by *SRSF2*^mut^-specific alternative splicing events (yellow = *SRSF2*^mut^ ASE, grey = normal transcript).

**Figure S9:**
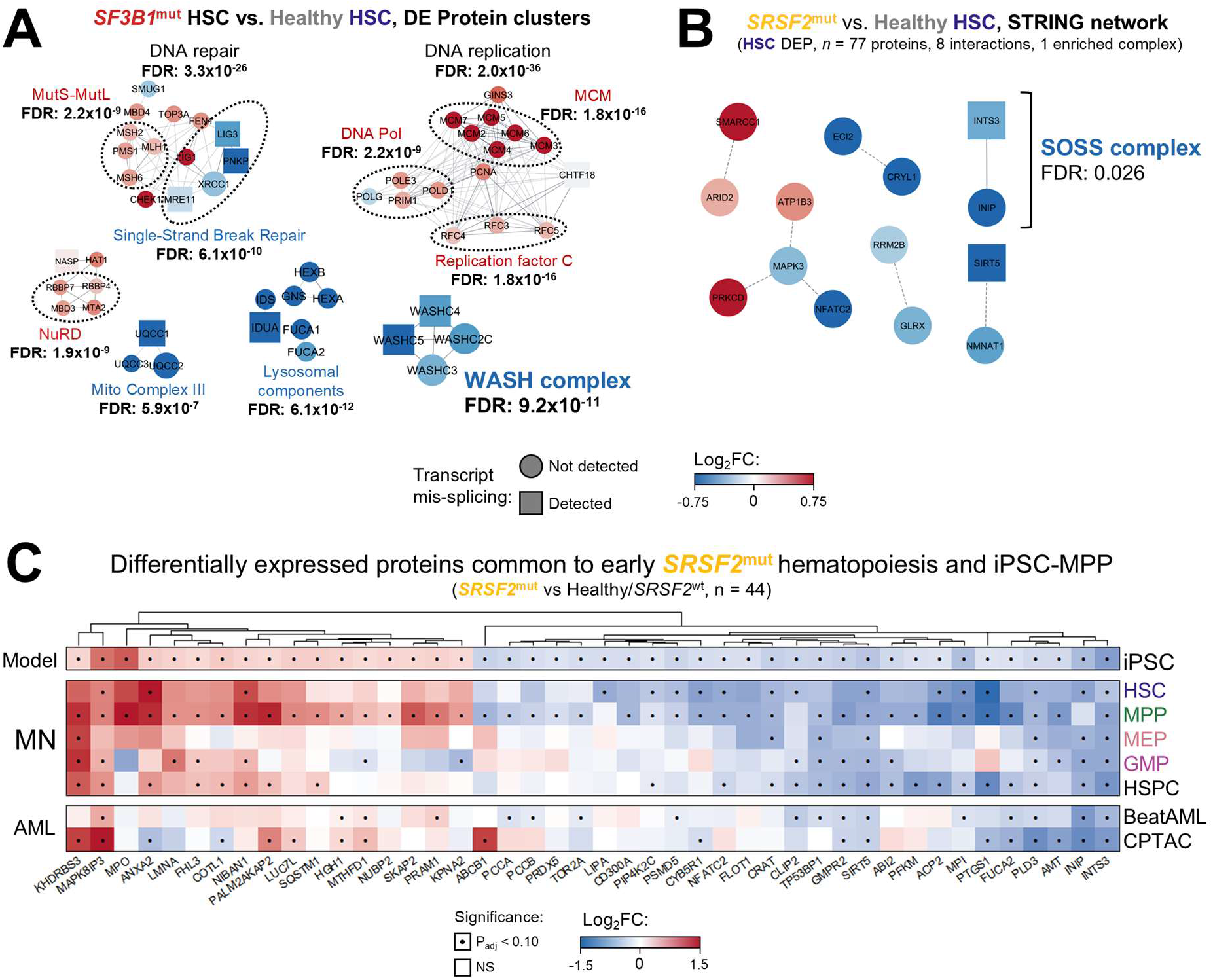
Proteomic phenotypes of *SF3B1*^mut^ and *SRSF2*^mut^ hematopoietic stem cells. A/B) Protein-protein interaction network of differentially expressed functional clusters in **a)** *SF3B1*^mut^ and **B)** *SRSF2*^mut^ hematopoietic stem cells. **C)** Relative protein heatmap of significantly differentially expressed proteins common between *SRSF2*^mut^ MN HSC/MPP and *SRSF2*^mut^ iPSC-derived MPP.

### Supplemental Methods

#### BM sample processing and sorting

Cryopreserved MNC were thawed in RPMI 1640 Glutamax (ThermoFisher) + 20% inactivated fetal bovine serum (FBS, ThermoFisher) + 100 U/mL DNase I (Sigma-Aldrich). BM MNC samples ≥ 20×10^6^ frozen cells were subjected to a magnetic-activated cell sorting (MACS) pre-enrichment step using anti-CD34 MicroBeads (Miltenyi Biotec) following the manufacturer’s protocol to facilitate later fluorescence-activated cell sorting (FACS) steps. Whenever pre-enrichment was performed, an unenriched aliquot of the thawed MNC was kept at 4°C for later MNC FACS-separation.

Following thawing (unenriched) or pre-enrichment (MACS), samples were labelled for FACS. Antibodies and fluorescent reagents used for sample labelling are listed in **Table S2**, including the respective RRIDs. Antibody labelling and washing steps were performed in (PBS + 2% FBS + 1 mM EDTA) kept at 4°C. Cells were analyzed and sorted according to downstream protocol requirements using a FACS ARIA II Fusion (Becton Dickinson) at the MedH FACS facility of Karolinska Institutet (sorting strategy in **Fig. S3a**). Fluorescence-minus-one (FMO) and single-stain controls were prepared on each experiment day. Data were analyzed with FlowJo v. 10.7.2 (Becton Dickinson).

#### Sample preparation and processing for 10X-ONT

Lin12^-^CD34^+^ live cells were viably purified onto tubes containing 10X Genomics resuspension buffer (PBS + 0.04% bovine serum albumin [Sigma]), left as single donor tubes or mixed at 50/50 male/female donor cell ratio for label-free sample multiplexing (**Figure S2A, Data S1**), and resuspended to a final concentration of 1,500 cells/µL.

All samples were loaded onto Chromium Single Cell Chips (10X Genomics) at a target capture rate of 15,000 cells per sample. Single-cell libraries were prepared using the Chromium GEM-X Single-cell 3’ v4 kit (10X Genomics), except full-length cDNA amplicons were generated from single-cell mRNA with a prolonged GEM-RT incubation time (2h instead of 45min), and amplified with increased incubation times (98°C 3 min, (98°C 15 sec, 63°C 20 sec, 72°C 3 min) *11 cycles, 72°C 3 min, 4°C ∞), so as to increase the resulting cDNA length.

Libraries were either pooled and sequenced on an Illumina NovaSeq 25B (Illumina) at 100 bp read length, or prepared individually for sequencing on a PromethION system (Oxford Nanopore Technologies) following the manufacturer protocol version SST_9198_v114_revO. For ONT, specifically, 10 ng of the cDNA amplicons were biotin-tagged and pre-amplified via a 4-cycle PCR using custom-ordered oligos and LongAmp Hot Start Taq 2X Master Mix (New England Biolabs). Biotinylated amplicons were subsequently isolated by pull-down utilizing M280 streptavidin magnetic beads (Invitrogen). Following bead purification, a post-pull-down PCR (4 cycles) was executed with PCR Primers (PRM; EXP-PCA001, listed in **Key Resources**) to amplify the target cDNA molecules. The amplified cDNA was purified with Agencourt AMPure XP beads (Beckman Coulter), and 200 fmol of the recovered sample was subjected to end-repair and dA-tailing using the NEBNext Ultra II End Repair/dA-Tailing Module (New England Biolabs). Sequencing adapters from the Ligation Sequencing Kit V14 (SQK-LSK114) were ligated to the end-prepped DNA using Salt-T4 DNA Ligase (New England Biolabs) and Ligation Buffer (LNB). The finalized libraries (50–100 fmol) were mixed with Sequencing Buffer (SB) and Library Beads (LIB) before being loaded onto an R10.4.1 PromethION Flow Cell (FLO-PRO114M). Data acquisition was performed with the MinKNOW software on a P2 solo device. Post run base-calling was performed using Dorado v.1.3.1 (Oxford Nanopore Technologies) in high-accuracy mode.

#### Sample preparation for pauciproteomics

Up to a maximum of 1000 cells from immunophenotypically-defined cell compartments (sorting strategy in **Fig. S3A**) were purified onto the center of dry 1.5 mL Protein LoBind (Eppendorf), snap frozen using dry ice and transferred to -80°C for storage until collection of each sample batch for mass spectrometry processing was complete.

Evotips PURE (EvoSep) were prepared in advance to receive samples by i) washing with 20µl of solvent B (0.1% formic acid in acetonitrile), centrifuging (60s, 800 g); ii) soaking in isopropanol for 1min; and iii) equilibrating with 20µl of solvent A (0.1% formic acid in water), centrifuging (60s, 800 g). The Evotips were then placed in an Evotip box containing 50mL of water to prevent drying.

Next, 5 µl of digestion buffer (0.2% DDM, 100 mM TEAB pH8.5, 20 ng/µl trypsin (Rapizyme MS grade, Waters)) was added to each sample (in ∼4µl of FACS buffer) in the original tube used in FACS sorting and mixed once by pipetting up and down. Samples were kept frozen on dry ice and only thawed immediately prior to addition of digestion buffer. The entire cell suspension mixed with digestion buffer (approx. 10 µl) was then transferred to a previously prepared Evotip. Evotips with samples were placed in the Evotip box containing 50mL of water and centrifuged for 3s. The Evotip box was then incubated at 37°C for 4 hours with mild shaking. After incubation, 50µl of Solvent A was added to the Evotips to wash and rinse the tips (centrifugation at 800 g, 60s). Finally, 200µl of Solvent A was added onto Evotips to also prevent drying from the top, as recommended by the manufacturer’s instructions.

Online LC-MS was performed using a Evosep One (Evosep, Odense, Denmark) coupled to a timsTOF SCP mass spectrometer (Bruker). The Evosep One LC system proceeded to inject sample onto the analytical column (Aurora Elite G3, C18, 15 cm long, 75 μm ID, 1.7 μm bead size, Ionoptics, Melbourne, Australia) constantly kept at 50°C by a heating oven (Column Toaster, Bruker, Bremen, Germany) using the Whisper 40 samples per day (40SPD) 31min gradient.

The timsTOF SCP mass spectrometer operated with the CaptiveSpray source, capillary voltage 1500 V, dry gas flow of 3 L/min, dry gas temperature at 200°C. Collision energy was set as 20 eV for 1/k0 0.60 V.s/cm and 59 eV for 1/k0 1.60 V.s/cm2. Data was acquired using Timscontrol v 6.0.6 and Compass HyStar 6.3.1.8. The above conditions were used for data independent acquisition (DIA) diaPASEF (parallel accumulation-serial fragmentation). Precursor ions were selected in the range of 100-1700 m/z and mobility range 0.64-1.45 1/k0 IM. MS/MS was used with Bruker default dia-PASEF settings with 1 MS1 ramp, 8 MS/MS ramps, 24 MS/MS windows, mass range was 400–1000 m/z, and mobility range was 0.64–1.37 V.s/cm2. Parallel accumulation and serial fragmentation were performed with a cycle time of 0.95 s.

#### RNA sequencing data analysis

##### Initial 10X-ONT data processing

Reads from both Illumina and Nanopore technologies were aligned against the GRCh38 reference genome. Illumina reads were processed using CellRanger v9.0.1 (10X Genomics). Nanopore reads were processed through the EPI2ME wf-single-cell workflow v3.3.0 (Oxford Nanopore Technologies) in 3prime:v4 mode with single nucleotide variant (SNV) calling enabled. Seurat v.5.4.0.^1^ was used to load, process and analyze the resulting data matrices.

##### Illumina-specific quality control and data processing

RNA count matrix filtering removed cells expressing less than 200 distinct genes, cells with a total percentage of mitochondrial reads above 8% and cells with a total UMI count below 2500 counts, as well as genes expressed in less than 3 cells. All samples were Log normalized, and the 2000 most highly variable genes across all samples were identified. These highly variable genes were then used to identify integration anchors and integrate all samples into one dataset. The healthy donor sample from batch 1 (of 3 Illumina batches) was used as the reference for sample integration. Louvain clustering was performed at a resolution of 1.8 to identify cells with similar gene expression profiles. The Seurat function *FindAllMarkers* was utilized to cluster marker genes, and these were subjected to EnrichR over-representation enrichment analysis with modules *GO_BP_2025*, *GO_MF_2025*, *CellMarker_2024*, *Azimuth_2023*, *Human_Gene_Atlas*, *Descartes_CTT_2021*, *HuBMAP_ASCTplusB_augmented_2022*, *PanglaoDB_Augmented_2021* and *CellMarker_Augmented_2021* to identify cluster cell type signatures. Clusters were manually labelled according to the above enrichment results. Pooled samples were demultiplexed into individual donor samples using souporcell.^2^

##### Nanopore-specific quality control and data processing

SNV genotyping matrix filtering removed cells with less than 100 distinct SNV genotypes, as well as SNV genotypes identified in less than 10 cells. Variable SNVs within each sample were identified and used to build dimensional reduction UMAPs, enabling the identification of major genotype clusters per sample. The Seurat function *FindMarkers* was utilized to identify SNVs differentially represented between genotype clusters. Per pooled sample, a minimum of 9 SNVs were utilized to distinguish both individuals (4 to 5 private SNVs per individual). Cells with either doublet genotypes (containing at least 2 of each set of private SNVs) or unidentifiable genotypes (containing none of the private SNVs) were removed from further analysis. RNA count matrix filtering removed cells expressing less than 200 distinct genes, cells with a total percentage of mitochondrial reads above 20% and cells with a total UMI count below 500 counts, as well as genes expressed in less than 3 cells. Downstream processing and annotation steps were performed as indicated above, except Louvain clustering was performed at a resolution of 3. Pooled samples were demultiplexed using the SNV genotype data together with gene expression data of Y-specific genes and the female-specific X chromosome gene *XIST*.

#### RNA expression analyses

##### 10X-ONT single-cell

For further analyses, the data were restricted to include only high-quality single cells that 1) passed QC in the separate data processing pipelines of both Illumina and Nanopore data processing and 2) had matching data on sex chromosome expression, souporcell-based Illumina sample demultiplexing and SNV-based Nanopore sample demultiplexing. Following both requirements resulted in a total number of 62,726 cells with short- and long-read sequencing information. Because of the higher read count and gene coverage per cell, differential RNA expression analyses were performed using Illumina short-read data. Cells were pseudobulked on a donor/cell type basis according to major hematopoietic progenitor lineages, and downstream gene expression analysis was performed with the use of DESeq2 v. 1.50.2^3^ (BH-corrected Wald test P-values). Pseudotime assignment was performed using Monocle3 v.1.4.26^4,5^, setting early multipotent cells (HSCMPP/HSCMPP-like) as the starting points for pseudotime determination.

##### Bulk

RNAseq data of MDS hematopoietic stem and progenitor cells were obtained from Shiozawa *et al*.^6^ and FASTQ read files aligned using STAR 2.7.10b.^7^ Mapped read pairs were counted through featureCounts v. 2.0.1^8^ and downstream gene expression analysis was performed with the use of DESeq2 v. 1.50.2^3^ (BH-corrected Wald test P-values).

##### RNA splicing analyses

A database of alternative splicing events [ASE] induced by the four splicing factor mutations *SF3B1*, *SRSF2*, *U2AF1* and *ZRSR2* was collated, based on the following studies:

| SF<sup>mut</sup> | Reference | PMID | Cell type |
| --- | --- | --- | --- |
| All four | Yoshida *et al.* Nature 2011<sup>9</sup> | 21909114 | MNC |
|  | Shiozawa *et al.* Nat Commun. 2018<sup>6</sup> | 30194306 | HSPC / MNC |
| SF3B1 | Dolatshad *et al.* Leukemia 2016<sup>10</sup> | 27211273 | HSPC |
|  | Bondu *et al.* Sci Transl Med. 2019<sup>11</sup> | 31292266 | MNC |
|  | Zhang J *et al.* Mol Cell 2019<sup>12</sup> | 31474574 | MNC |
|  | Clough *et al.* Blood 2022<sup>13</sup> | 34861039 | iPSC |
|  | Asimomitis *et al.* Blood Adv. 2022<sup>14</sup> | 35042235 | iPSC |
|  | Moura *et al.* Cancer Res. 2024<sup>15</sup> | 37921711 | HSPC and erythroid cells |
|  | Moura *et al.* Blood Neoplasia 2024<sup>16</sup> | 40454400 | Hematopoietic stem cells |
|  | Sarchi *et al.* Blood Cancer Discov. 2024<sup>17</sup> | 38856693 | HSPC |
|  | Rombaut *et al.* Nature Commun. 2024<sup>18</sup> | 38589367 | MNC |
| SRSF2 | Yoshimi *et al.* Nature 2019<sup>19</sup> | 31578525 | HSPC / MNC / AML |
| U2AF1 | Jin *et al.* Cancer Res. 2024<sup>20</sup> | 38417135 | AML |
| ZRSR2 | Madan *et al.* Nat Commun. 2015<sup>21</sup> | 25586593 | MNC |

A re-analysis of differential splicing within the cited bulk RNAseq data of Shiozawa *et al*.^6^ was performed using rMATS v. 4.1.1^22^ (BH-adjusted likelihood-ratio test P-values). Cut-offs for statistically significant differentially spliced junctions were False Discovery Rate < 0.001, minimum average count > 10, absolute percent spliced-in difference (|ΔPSI|) > 0.10. Together, these data were utilized to map junctions of interest and retrieve junction count PSIs within individual cells of 10X-ONT long-read data. 10X-ONT per-cell PSIs were either overlaid on Illumina data, pseudobulked across cell lineages for statistical comparison or binned across Monocle3 pseudotime to identify dynamics over cell differentiation. Pseudobulked PSIs were compared through non-parametric Wilcoxon rank-sum tests of SF^mut^ PSIs against all other SF^mut^ as well as between each SF^mut^ pair. Sashimi plots for ASE visualization were generated using ggsashimi v. 1.1.5.^23^

Additionally, since the total number of *SF3B1*^mut^ and *ZRSR2*^mut^ ASEs numbers in the hundreds, we verified the corresponding alternatively spliced junctions through manual analysis and visualization in our own data as well as three of the cited studies, and could reliably trace full isoform characteristics of *SF3B1*^mut^ ASEs within 10X-ONT data (summarized in **Fig. 2** and **Fig. S4**). By contrast, since the reported *SRSF2*^mut^ and *U2AF1*^mut^ ASEs number in the thousands within the above studies alone, we instead utilized nonsense-mediated decay (NMD) information from Yoshimi *et al*.^19^ to assess NMD/protein effects within the pauciproteomics cohort. In line with our *SF3B1*^mut^ data, most of the reported *SRSF2*^mut^ NMD effects did not have an actual discernible consequence either in transcriptomics (in the cited studies or in our own 10X-ONT data), or proteomics.

#### Pauciproteomics data analysis

##### Statistical analysis of proteomic intensities

Raw mass spectrometry data files were analyzed using Spectronaut vs. 19.4 (Biognosys) with the DirectDIA workflow for label-free quantification. Data search parameters included 1% FDR (false discovery rate) on both protein and peptide level, 2 as maximum missed cleavages, and Trypsin as enzyme type and cleavage rules. The search was performed against the Human Swissprot protein database (2024-01-10, 20413 canonical protein entries). The resulting MS2 protein intensity matrices from Spectronaut analysis were transformed using the variance-stabilizing normalization (VSN) function *vsn2* from the R package *vsn*,^24^ and batch-corrected per mass spectrometry run batch (*n* = 3 batches) using the *removeBatchEffect* function from *limma*.^25^ Proteins where insufficient detection was achieved were excluded from the dataset (detected in <50% of samples from either group under comparison). Following these initial steps, statistical comparisons of mass spectrometry data were performed by pairwise comparison using Benjamini-Hochberg (BH) multiple comparison-corrected unpaired *t*-tests (comparing intensities) and Fisher’s exact tests (comparing detection ability).

##### Protein network clustering and module identification

A human protein–protein interaction (PPI) network was built using the STRING v12.0 database^26^ (protein.links.v12.0.txt), with genes as nodes and interactions as edges. Interactions were restricted to a minimum score of 700. Differentially expressed genes associated with each SF^mut^ group (SF3B1 and SRSF2) and data modality (10X-ONT, pauciproteomics) were intersected with the nodes. The resulting subgraph was extracted for downstream module detection. Candidate modules were seeded by exhaustive enumeration of maximal cliques, retaining cliques of size ≥ 3 nodes. Because maximal-clique enumeration produces redundancy among overlapping cliques, an iterative merging procedure was applied to resolve highly overlapping cliques into overarching modules. At each iteration, pairwise Jaccard similarity values were computed between all cliques based on shared member genes, and clique pairs with Jaccard similarity ≥ 0.5 were connected in an auxiliary clique-overlap graph. Connected components were identified within this auxiliary graph, and subjected to Leiden clustering^27^ (modularity-objective function, default settings) for partitioning overlapping cliques into communities. Cliques within the same Leiden community were merged by taking the union of their member genes to form a new candidate module. Cliques not involved in any high-overlap pair were carried forward unchanged. This procedure was repeated until no clique pairs retained a Jaccard similarity value of ≥ 0.5. Because individual proteins could remain assigned to more than one module after convergence, the mean STRING score between multiply assignable proteins and all other members of each candidate module was computed, and the protein was assigned to the module with the highest mean score. Following this reassignment, modules reduced to a single member were merged into one of the original multi-member modules, again using the mean STRING score to select the best-matching module. Singleton modules with no scoring evidence for merging were discarded.

##### Functional annotation of protein modules

For each module identified above, over-representation analysis was performed using protein complexes and Gene Ontology (GO) gene sets. Reference protein complex annotations were retrieved from OmnipathR, which aggregates complex membership from multiple source databases.^28^ Complexes with fewer than two member genes were excluded from this enrichment analysis. GO gene sets (combined Biological Process, Cellular Component, and Molecular Function categories) were obtained from the MSigDB C5 collection (v. 2026.1)^4^. Enrichment was assessed at a Benjamini–Hochberg-adjusted p-value cutoff of 0.01, with a minimum of two overlapping genes required for a term to be retained. To reduce redundancy between adjacent modules with highly similar functional signatures, pairwise Jaccard similarities of enriched GO term identifiers were computed between all module pairs, and pairs with similarity ≥ 0.5 were merged. After this enrichment analysis, cliques of 2 nodes were split into singletons, and singletons were assigned to the top-scoring protein module, if enriched in a minimum of 5 protein subcomplexes related to the module. Finally, cliques were filtered to include only clusters with described protein complex identity as well as enrichment within the Gene Ontology database.

#### Tandem Mass Tag proteomics data analysis

Analysis of Tandem Mass Tag (TMT) data from external proteomic cohorts was pursued to validate the findings of pauciproteomics within independent cancer cohort datasets including SF^mut^ patients. All cohorts had been deposited with open public access within the NCI’s Proteomic Data Commons database.^29^ In each case, pre-normalized semi-quantitative data resulting from the PDC’s Common Data Analysis Pipeline were annotated and compared through non-parametric Wilcoxon rank-sum tests. Genomic data were sourced from the publications noted below as well as mapped by cBioPortal.^30–32^ *BeatAML 1.0*:^33–35^ After quality control, this cohort includes 9 *SF3B1*^mut^ patients, 16 *SRSF2*^mut^ patients, 2 *U2AF1*^Q157^ patients and 1 *U2AF1*^S^^34^ patient.

*CPTAC AML*:^36^ Only one sample source was analyzed per patient, with preference to bone marrow material. After quality control, this cohort includes 6 SF3B1^mut^ patients and 14 SRSF2^mut^ patients. *CPTAC Pan-Cancer*:^37,38^ Across the CPTAC Pan-Cancer cohort, 22 tumors were reported as *SF3B1*^mut^. These were further filtered to include only cases with VAF > 15% and carrying hotspot mutations, resulting in 6 high-confidence cases (2 K700E, 1 T663I, 1 V701F, 1 K666N, 1 G740E). Tumor expression data was therefore restricted to clear cell renal cell carcinoma (CCRCC), non-clear cell renal cell carcinoma (non-CCRCC), lung adenocarcinoma (LUAD_Disc, LUAD_Conf) and breast cancer (BC_Prospective) and normalized against the mean normalized protein expression of each tumor type.

### Supplemental tables

**Table S1:** List of genes profiled in each myeloid neoplasm patient cohort.

| Genes profiled by all three NGS panels |  |  |  |  |  |  |
| --- | --- | --- | --- | --- | --- | --- |
| *ABL1* | *CEBPA* | *EZH2* | *JAK3* | *NRAS* | *SAMD9* | *STAG2* |
| *ASXL1* | *CREBBP* | *FLT3* | *KDM6A* | *PDGFRA* | *SAMD9L* | *STAT3* |
| *BCL2* | *CSF3R* | *GATA1* | *KIT* | *PHF6* | *SETBP1* | *TET2* |
| *BCOR* | *CUX1* | *GATA2* | *KRAS* | *PPM1D* | *SETD2* | *TERT* |
| *BCORL1* | *DDX41* | *GNB1* | *MPL* | *PRPF8* | *SF3B1* | *TP53* |
| *BRAF* | *DNMT3A* | *IDH1* | *MYD88* | *PTEN* | *SH2B3* | *U2AF1* |
| *CALR* | *EP300* | *IDH2* | *NF1* | *PTPN11* | *SMC1A* | *WT1* |
| *CBL* | *ETNK1* | *JAK1* | *NOTCH1* | *RAD21* | *SMC3* | *ZRSR2* |
| *CDKN2A* | *ETV6* | *JAK2* | *NPM1* | *RUNX1* | *SRSF2* |  |
| Genes profiled by two of three NGS panels |  |  |  |  |  |  |
| *AKT1* | *CBLB* | *DICER1* | *IKZF1* | *MYC* | *ROBO1* | *U2AF2* |
| *ALK* | *CCND3* | *DNM2* | *IL7R* | *NF2* | *ROBO2* | *UBA1* |
| *ANKRD26* | *CDK4* | *EED* | *IRF1* | *NFE2* | *RPL5* | *USP9X* |
| *ARHGEF10* | *CDKN2B* | *EIF6* | *JARID2* | *NIPBL* | *RRAS* | *ZBTB7A* |
| *ARID1A* | *CHEK2* | *EGFR* | *KMT2A* | *NOTCH2* | *SF1* | *ZEB2* |
| *ARID2* | *CSF1R* | *FBXW7* | *KMT2C* | *NXF1* | *SF3A1* |  |
| *ASXL2* | *CSF2RB* | *FGFR2* | *KMT2D* | *PHIP* | *SMARCA4* |  |
| *ATRX* | *CSNK1A1* | *GFI1* | *LUC7L2* | *PIK3CA* | *SRP72* |  |
| *BAP1* | *CTCF* | *GIGYF2* | *MED12* | *RAD51* | *STAG1* |  |
| *BCL11B* | *DCC* | *GNAS* | *MGA* | *RB1* | *STAT5B* |  |
| *BRCC3* | *DHX15* | *HRAS* | *MYB* | *RIT1* | *SUZ12* |  |

**Table S2:**
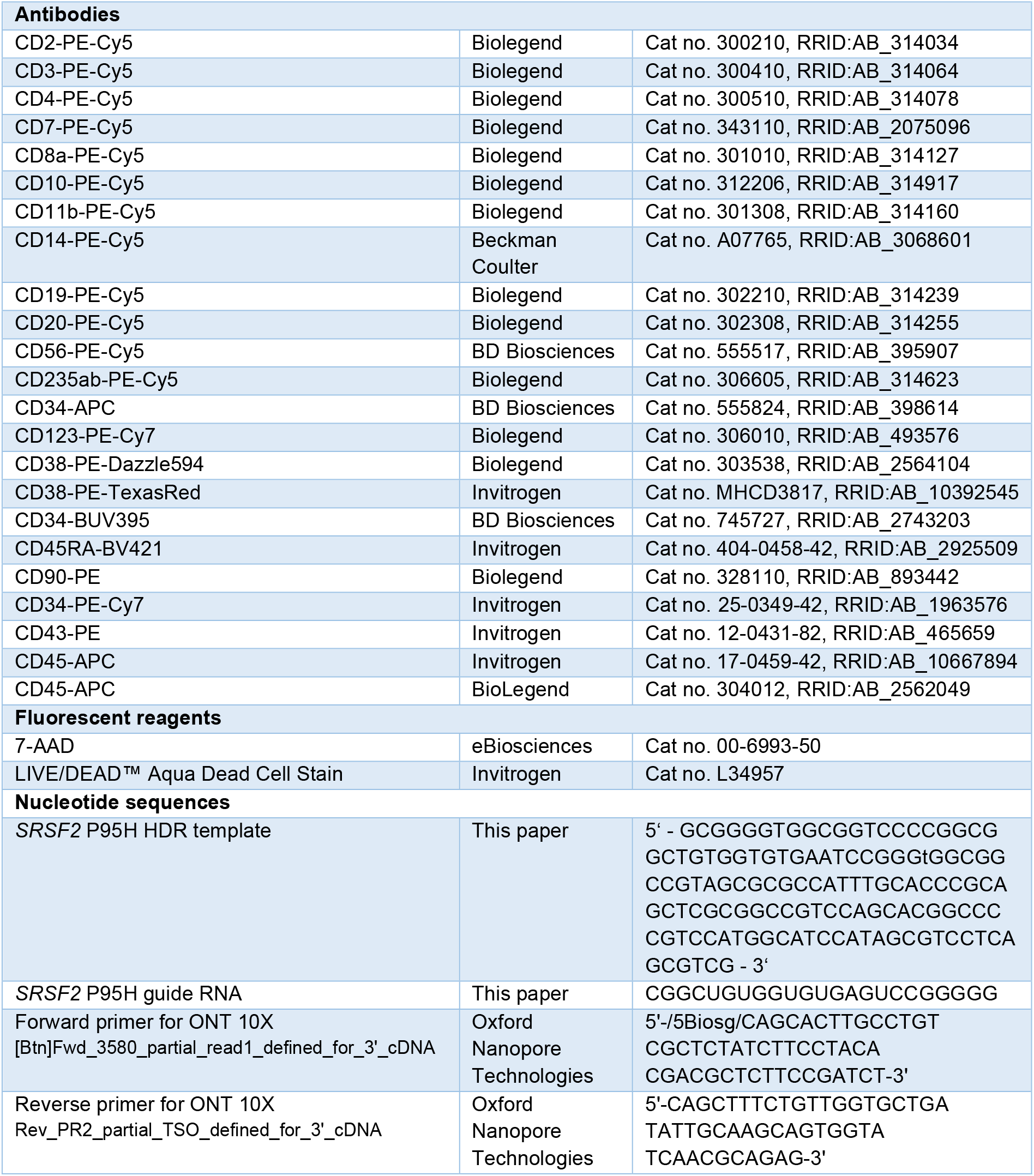
Key resources.

